# netPCF: Geometry-aware pair correlation functions for spatial biology

**DOI:** 10.64898/2026.07.02.736020

**Authors:** Joshua W. Moore, Joshua A. Bull, Helen M. Byrne

## Abstract

Spatial organisation is a defining feature of biological systems, underpinning cellular interactions, tissue function, disease progression and therapeutic response. Identifying and quantifying spatial organisation may require methods that resolve relationships across spatial scales. The pair correlation function (PCF) quantifies spatial dependence between points across multiple length scales, but its standard Euclidean formulation is poorly suited to data defined on irregular, curved or otherwise structured domains, where tissue geometry may constrain biological organisation and distort Euclidean distances. Here, we introduce netPCF, a geometry-aware extension of the PCF for quantifying spatial organisation on complex biological domains. By representing tissue structures, anatomical surfaces and other constrained geometries as spatial networks, netPCF generalises the PCF beyond extrinsic Euclidean settings. The framework derives the expected behaviour of the statistic under complete spatial randomness using interpretable finite-support kernels, provides bootstrap-based uncertainty quantification, and includes practical criteria for assessing domain discretisation adequacy. We further extend netPCF to marked (labelled) biological data using feature kernels for categorical and continuous attributes, enabling unified analysis of cell identities, marker intensities, phenotypic states, gene expression and other quantitative features on structured domains in any spatial dimension. All methods are implemented in the open-source Python package `spacenet`.

Synthetic studies show that netPCF recovers classical Euclidean behaviour on sufficiently resolved networks and is robust to common imaging noise. We demonstrate its utility in two biological applications. In three-dimensional imaging mass cytometry data from HER2+ breast carcinoma, netPCF separates tissue architecture-driven proximity from biologically meaningful endothelial and immune cell organisation. In reconstructed surfaces of developing murine embryos, netPCF identifies a transition in the *Wnt1* –*Wnt6* relationship from short-range co-localisation at E9.5 to spatial exclusion at E11.5, a pattern of ectodermal boundary refinement not captured by prior voxel-wise co-expression analysis. Overall, netPCF provides a statistically grounded and practical framework for quantifying spatial organisation on complex biological domains.

**Author summary:** Spatial organisation is central to many biological processes, but it is often measured using distances that ignore the shape of the tissue or structure being studied. We introduce netPCF, a method for quantifying multiscale spatial correlation in data that lie on complex biological domains, including irregular, curved, or branching structures. netPCF reconstructs the domain as a distance-preserving spatial network and estimates pair correlation along this intrinsic geometry, allowing spatial associations to be interpreted relative to the structure in which they occur. The framework includes uncertainty estimates and extensions for categorical and continuous markers, supporting analysis of cell types, marker intensities, phenotypic states, and gene expression patterns. In synthetic data, netPCF recovers expected spatial behaviour on well-resolved networks. In biological imaging data, it distinguishes apparent cell proximity caused by breast carcinoma tissue architecture from biologically meaningful cell organisation, and reveals a developmental transition in Wnt gene organisation over the surface of a murine embryo that direct co-expression analysis does not capture. netPCF is available in the open-source Python package `spacenet`, supplemented with online tutorials supporting practical use across spatial biology applications.

## Introduction

The pair correlation function (PCF), also known as the radial distribution function, is widely used to quantify spatial relationships between points across multiple length scales. Originally developed in statistical physics to study particle systems [51], the PCF is now used across ecology, astronomy, materials science, and spatial biology [27, 46, 58, 14]. Its central appeal is that it measures spatial dependence as a function of length scale, distinguishing between clustered, random, and regular point patterns. In biomedical image analysis, this scale-resolved perspective is valuable because tissue organisation is rarely captured by nearest-neighbour distance or adjacency relation [15]. PCFs and related second-order statistics have therefore been used to quantify cell–cell interactions [37], tumour microenvironment organisation [41], immune infiltration [59, 12], and morphogenetic patterning in developmental systems [45].

A persistent challenge in applying PCFs to biological systems is that spatial data is rarely embedded in simple Euclidean domains. Cells, molecules and tissue features are often constrained by the geometry of the biological domain they occupy, including epithelial boundaries, stromal compartments, vascular structures, luminal spaces, curved surfaces, and branching architectures [36, 31, 52, 54, 42] (Fig. 1). In these settings, a global notion of Euclidean distance may not reflect the spatial relationships available within the biological domain. Apparent clustering can arise because two populations are restricted to the same anatomical compartment, while spatial apparent exclusion can arise because regions of the domain are not contiguous. Thus, ignoring the intrinsic geometry of the domain of interest can obscure or distort the spatial correlations that PCFs are intended to measure [30].

**Fig 1.**
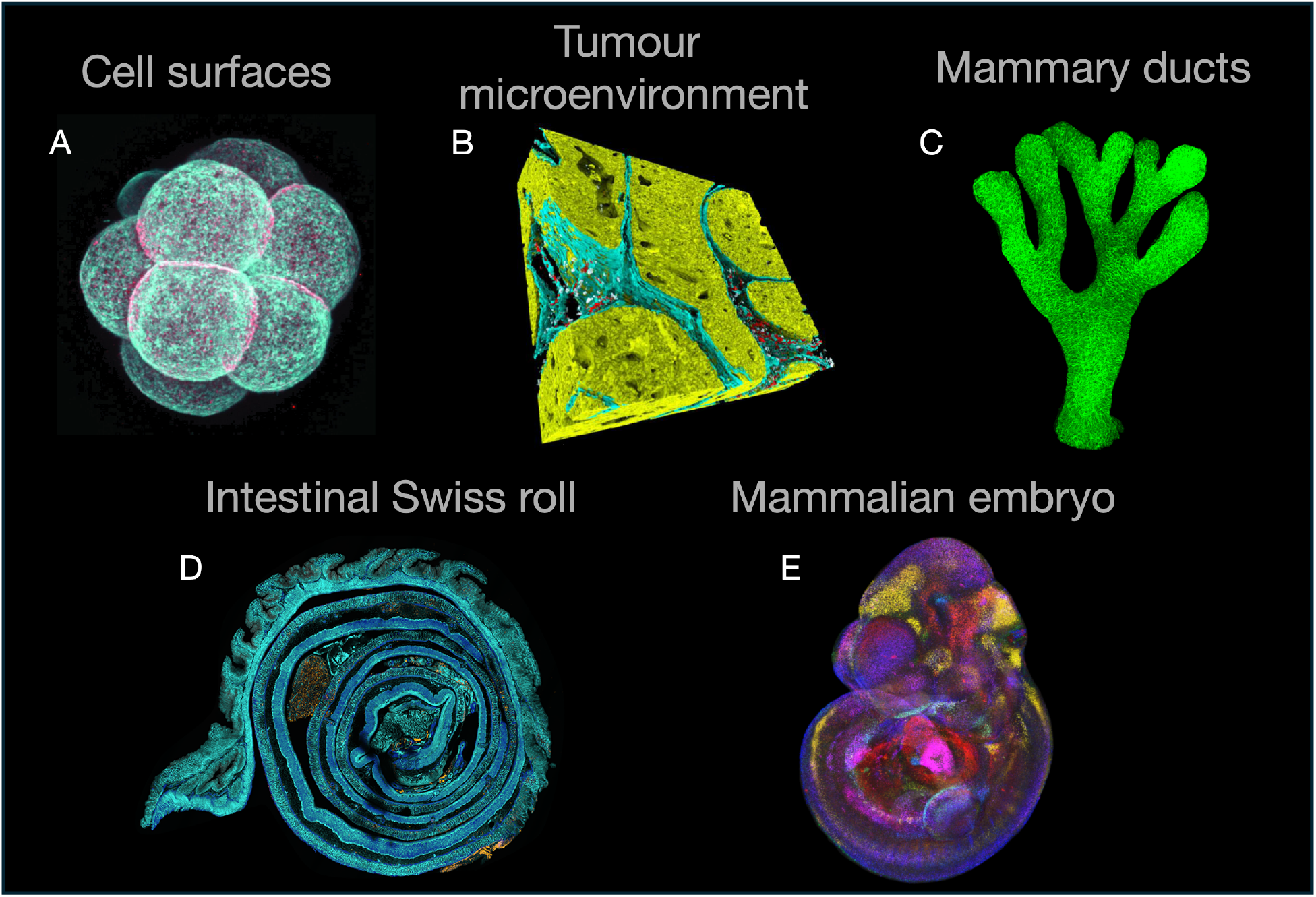
Complex geometries in spatial biology. Examples of biological imaging with nontrivial underlying geometry. (A) Fluorescent confocal imaging of an mTmG (blue) and LifeAct-GFP (red) transgenic murine embryo during the 8-cell stage, reproduced from [36]. (B) Three-dimensional imaging mass cytometry of HER2+ breast carcinoma, showing cells expressing SMA (blue), vWF/CD31 (red), panCK (yellow), and CD8*α* (white), reproduced from [31]. (C) Confocal maximum-intensity projection of an E17.5 wild-type mouse mammary gland whole-mount stained for the epithelial marker EpCAM (green), reproduced from [52]. (D) Swiss-roll preparation of a 2-month-old mouse large intestine, with dividing cells highlighted in yellow (Ki67), cell membranes in turquoise, and fibroblasts and muscle cells in blue, reproduced from [54]. (E) Spatial gene expression patterns of the Wnt and Fzd families in an E11.5 murine embryo captured by *in situ* hybridisation and optical projection tomography, reproduced from [42].

One strategy for representing complex spatial organisation is to use local geometric information to approximate global structure. Methods based on this principle construct discretised representations of space in which local relationships encode the underlying geometry. For example, meshes built from local connectivity rules can capture tissue microstructure [38, 53], while persistent homology methods, such as alpha filtrations, can recover approximations of tissue surfaces and other manifold-like structures [62, 29]. Similarly, triangulation-based and graph-based reconstruction approaches represent spatial organisation across multiple resolutions [17, 57].

These methods can be viewed collectively as spatial networks: graphs whose nodes are embedded in space and whose edges encode local connectivity. When edge weights reflect physical or geodesic distance, such networks provide an approximate representation of the underlying spatial domain. Despite their methodological differences, these constructions share the same underlying idea: complex geometry can be represented through local interactions that collectively approximate global structure [4].

Pair correlation functions have previously been adapted to be applied to network topologies. Okabe and colleagues extended Ripley’s *K*-function, a cumulative analogue of the PCF, to linear networks [47], followed by related developments for PCFs on network domains [3]. Building on this work, Baddeley and co-workers introduced geometric correction factors for *K*-functions and PCFs on networks, ensuring that these estimators are well defined and account for the amount of network accessible at each distance [1]. Subsequent work improved computational efficiency, including fast Fourier transform approaches for accelerating network-based estimation [50]. These methods provide a rigorous foundation for spatial statistics on networks such as roads, rivers, and transport systems, where the network is fixed and known in advance. Recently, attention has focused on the relaxing the requirment for prescribed networks for correlation estimations on complex domains. For example, PCFs applied to Voronoi topology [60] or regular spatial discretisation [30, 25] have been considered but without preserving the spatial resolution.

In biomedical imaging, the spatial support is often unknown, irregular, and must be inferred directly from the data. The network is therefore not merely a pre-existing domain on which points are observed; it is a data-driven approximation to the biological structure being analysed. This raises two related challenges. First, the method must preserve biologically meaningful spatial scale by measuring distances along the inferred domain, rather than only representing topological adjacency. Second, it must avoid introducing artefacts from the domain discretisation. Existing Euclidean PCFs ignore intrinsic tissue geometry [30], while existing network PCFs do not address the case where the network is constructed from biological point clouds to approximate the spatial domain. At present, there is no accessible and practical PCF framework that jointly supports data-driven spatial networks, distance-preserving domain approximation, and marked biological measurements.

In response, we introduce the network-based pair correlation function (netPCF), an extension of the PCF for data-driven spatial networks that approximate biological structures. netPCF measures pairwise distances along an inferred spatial network and applies a geometric correction based on the accessible network length at each radius. This enables scale-dependent spatial association to be estimated on arbitrary spatial graphs, including curved, branching, perforated, surface, and volumetric domains. To quantify uncertainty and spatial heterogeneity in these correlation estimates, we adapt bootstrap methods from spatial statistics for arbitrary spatial networks using a novel volume-balanced partition algorithm, enabling local variability in spatial correlation to be assessed. We also derive practical conditions for sufficient kernel coverage, providing data-dependent criteria to ensure that discretisation of the spatial network does not bias the resulting correlation estimates.

We further extend netPCF using feature kernels, so that categorical and continuous node attributes can be analysed within the same framework. This is important for spatial biology, where nodes commonly represent image pixels, cells or tissue objects annotated with marker intensities, phenotypes, transcript counts, or morphological measurements. By changing only the feature kernels, the same network-based formulation recovers unmarked netPCFs, cross-netPCFs for categorical cell types, and weighted netPCFs for continuous molecular or morphological features. Thus, netPCF provides a unified approach for quantifying marked spatial organisation independently of the geometry or spatial dimension of the data. These methods can be accessed via the publicly available Python package `spacenet` and is documented at https://www.spacenet-python.com.

We validate netPCF by showing that, on sufficiently resolved Delaunay networks, it recovers classical Euclidean PCF behaviour for point processes with known second-order structure. We then demonstrate that Euclidean PCFs can be systematically biased by geometric constraints, such as excluded regions, branching domains, and curved structures, whereas netPCF recovers the expected null behaviour by measuring distances along the intrinsic domain geometry. We evaluate robustness to practical sources of uncertainty in imaging-derived spatial networks, including reduced sampling density, edge-length filtering choices, and positional noise.

We demonstrate the utility of netPCF through the re-analysis of two independent three-dimensional imaging datasets. The first is imaging mass cytometry data from HER2+ breast carcinoma [31]. This case study shows that geometry-aware pair correlation analysis can alter the biological interpretation of spatial proximity: apparent endothelial–immune co-localisation in the full tissue domain is largely explained by tumour architecture, whereas immune–immune aggregation, including B cell–CD8+ T cell co-localisation, persists after restricting the analysis to the tumour microenvironment. These results illustrate how netPCF can distinguish geometry-driven proximity from intrinsic cellular organisation in complex biological tissues.

The second re-analysis focuses on Wnt gene expression over the surface of the developing murine embryo [42]. By analysing spatial correlations among *Wnt1, Wnt2*, and *Wnt6* at E9.5 and E11.5, we show that voxel-wise co-expression alone is insufficient to capture multiscale reorganisation of gene expression domains over the curved embryonic surface. In particular, netPCF reveals a developmental transition in the *Wnt1* –*Wnt6* relationship, from short-range surface association at E9.5 to spatial exclusion at E11.5, despite previous analyses reporting no convincing voxel-level co-localisation of *Wnt6* with other Wnt genes. Together, these examples demonstrate that netPCF provides a general framework for estimating spatial correlation in complex biological domains, enabling biological interactions to be interpreted in their intrinsic geometric context.

## Methods

### Inferring Spatial Networks from Spatial Data

Let 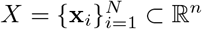 denote a set of observed spatial locations extracted from spatial data. We represent these data as a spatial network *G* = (*V, E*), where each node *v*_*i*_ ∈ *V* corresponds to an observation at location **x**_*i*_, and an edge *e*_*ij*_ ∈ *E* encodes a local neighbourhood relationship between points *i* and *j*. In our setting, nodes may represent any spatially embedded entity, for example, cell centroids extracted from a segmentation mask or image pixels.

To construct the network, we use the Delaunay triangulation of the point set. This provides a parameter-free way to define local connectivity by linking nearby points while avoiding degenerate simplices and overly dense neighbourhoods. Under sufficiently dense sampling, the resulting triangulation yields a piecewise linear approximation to the underlying spatial domain, so that local graph structure reflects local geometry [4]. Each edge is assigned the Euclidean weight, *w*_*ij*_ = ∥**x**_*i*_ − **x**_*j*_∥_2_, which measures the physical separation between the corresponding nodes. When the sampling density is sufficiently high, these local Euclidean edge lengths provide a discrete approximation to intrinsic distances on the domain.

To control the scale of connectivity and suppress spurious long-range links, we optionally apply an edge-length threshold and retain only edges satisfying *w*_*ij*_ *< ϵ*. This filtering step preserves the local structure of the triangulation while restricting the graph to interactions occurring over short spatial ranges. Distances between nodes are then defined using the weighted shortest-path metric on the resulting graph, *d*(*v*_*i*_, *v*_*j*_) = *d*_sp_(*v*_*i*_, *v*_*j*_). This metric approximates geodesic distance along the reconstructed spatial domain by concatenating local Euclidean edge segments, rather than measuring straight-line distance through the ambient space. Herein, let *d*_sp_(*v*_*i*_, *v*_*j*_) = *d*_*ij*_. Similar graph-based distance constructions are standard in manifold learning and spatial statistics, and provide a natural way to encode intrinsic geometry in discrete data.

In addition, each node may have associated attributes, such as categorical labels or continuous measurements (or both). We denote 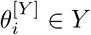, for *Y* the space of attributes. The set, *Y* could represent a discrete set of categories (e.g. cell types annotations), or continuous values (e.g. proteomics intensity or transcript counts). This network representation provides a geometry-aware discretisation of the spatial domain that supports estimation of spatial statistics using distances defined on the reconstructed structure.

An example of domain reconstruction using a Delaunay network is shown in Fig. 2, where the relationship between Euclidean edge weights and weighted shortest-path distances is illustrated.

**Fig 2.**
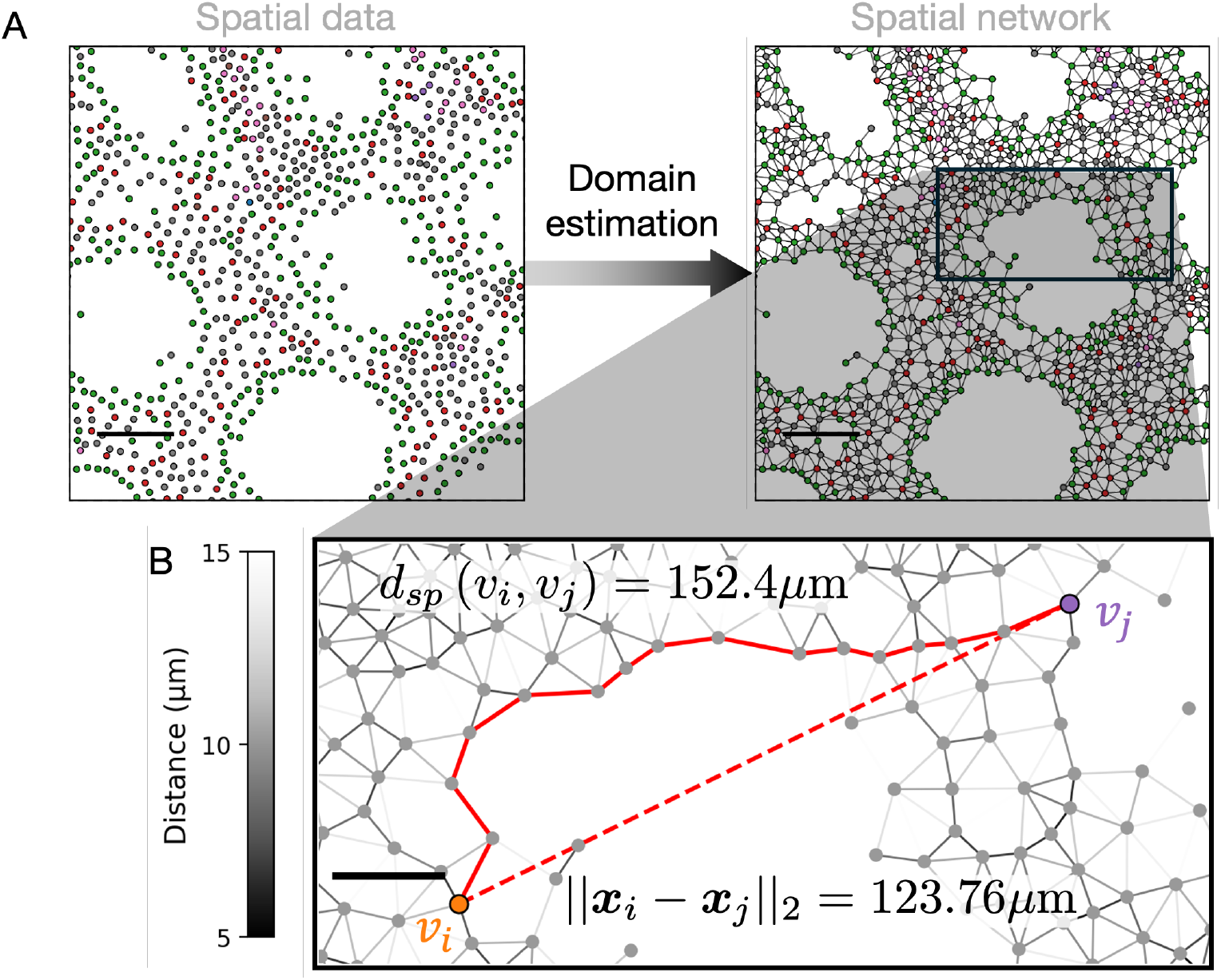
Spatial domain estimation using weighted Delaunay networks. (A) Schematic illustrating the reconstruction of a spatial domain from point data within a two-dimensional 300 × 300 µm^2^ region using a Delaunay triangulation with an edge-length filter of 15. Scale bar, 50 µm. (B) Magnified view of the resulting spatial network highlighting edge weights corresponding to Euclidean distances (greyscale). The distance-weighted shortest-path distance and the direct Euclidean distance between nodes *v*_*i*_ and *v*_*j*_ are indicated by the solid red and dashed lines, respectively. Scale bar, 20 µm.

### Estimation of Pair Correlation Functions on Spatial Networks

The estimated pair correlation function (PCF) quantifies spatial dependence by comparing the observed density of point pairs at separation *r* to the expected density under a specified null model,

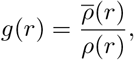

as discussed in [3]. We take the homogeneous Poisson point process, corresponding to Complete Spatial Randomness (CSR), as the null model (although alternative null models can be defined for point patterns on networks, their derivations remains an active area of research). We use CSR as a practical and interpretable baseline for assessing spatial structure in biological data. In this reference, at a distance *r >* 0, values *g*(*r*) *>* 1 indicate clustering, values *g*(*r*) *<* 1 indicate inhibition, and values *g*(*r*) ≈ 1 are consistent with spatial randomness [3, 13].

To accommodate non-Euclidean geometry, we define the PCF on a spatial network *G* = (*V, E*) by replacing Euclidean distances with the weighted shortest-path distance *d*_*ij*_ between nodes *v*_*i*_ and *v*_*j*_. This yields a network-based pair correlation function (netPCF) of the form

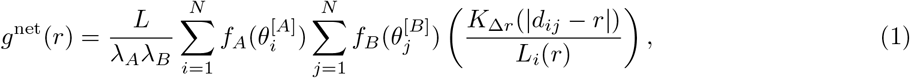

where *L* = ∑_*i,j*_ *w*_*ij*_ denotes the weighted total length of the network domain, *f*_*A*_ : *A* → ℝ and *f*_*B*_ : *B* → ℝ are feature functions used to select or weight nodes according to categorical or continuous attributes, associated with the node feature space *A* and *B*, respectively (for explicit forms of feature functions, see Supplementary Information 1.3). The corresponding effective feature intensities are

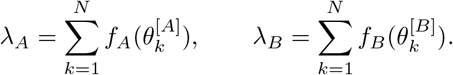

The term *L*_*i*_(*r*) accounts for the accessible network length at distance *r* from node *v*_*i*_ and ensures appropriate normalisation under CSR:

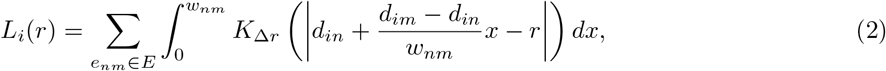

where *w*_*nm*_ is the Euclidean length of edge *e*_*nm*_ ∈ *E* and *K*_Δ*r*_ is a finite-support spatial kernel. In this study, we consider polynomial kernels of the form

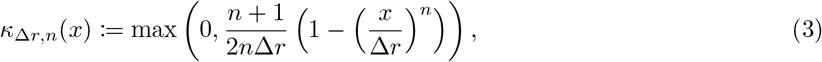

for the kernel shape parameter *n* ∈ ℝ_*>*0_ and finite support bandwidth Δ*r* ∈ ℝ_*>*0_. We then take *K*_Δ*r*_(*x*) = *κ*_Δ*r,n*_(*x*) throughout. This kernel provides smooth, compactly supported weighting over distance intervals, with the relative emphasis placed on contributions near the target distance controlled by the choice of *n*. Moreover, allowing the kernel shape parameter to vary provides a continuous family of weighting functions, which helps bridge the discrete network representation and the underlying continuous spatial domain. Common kernels used in spatial statistics can be constructed with Eq. (3), including the triangular kernel for *n* = 1, the Epanechnikov kernel for *n* = 2, and the top-hat (uniform) kernel in the limit as *n* → ∞ [3] (see Supplementary Information 1.1 for further details.) Using Eq. (3), an explicit expression for *L*_*i*_(*r*) can be derived, enabling efficient computation (see Supplementary Information 1.2 for details). An example of the polynomial kernel and its associated normalisation over a network is shown in Fig. 3.

**Fig 3.**
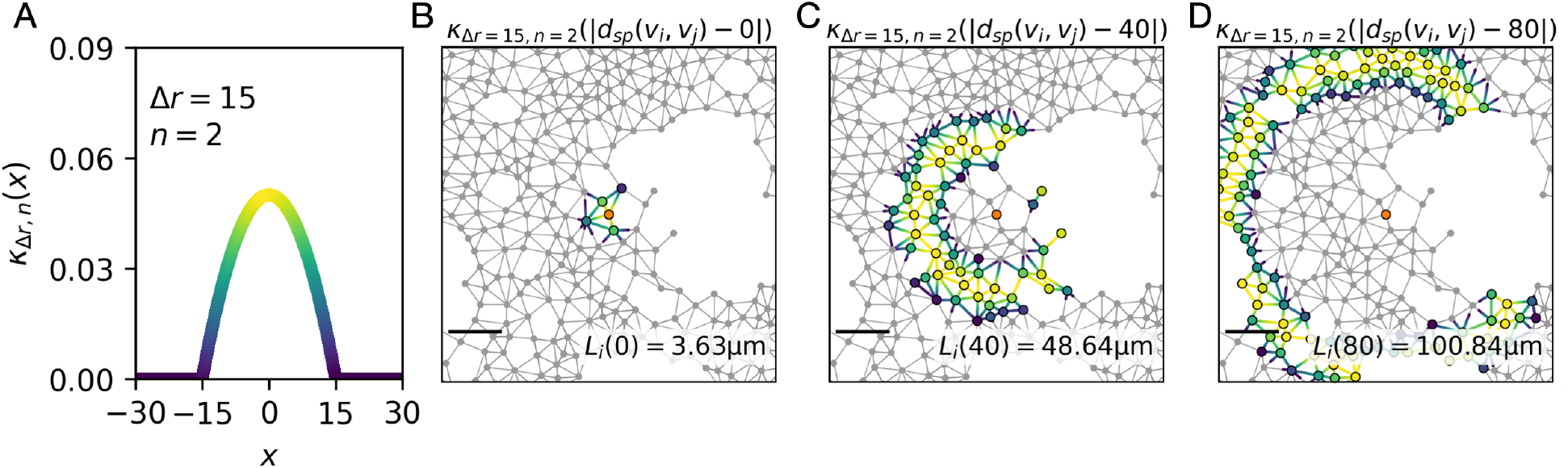
Spatially resolved polynomial kernel on a network. (A) The polynomial kernel of finite support (Eq. (3)) is shown with parameters *n* = 2 and Δ*r* = 15, evaluated around an exemplar node *v*_*i*_ (highlighted in orange) at distances *r* = 0, 40, and 80 µm (B-D). The corresponding normalisation term, *L*_*i*_(*r*), representing the total kernel volume over the network centred at node *v*_*i*_, is also illustrated.

Overall, the estimator (Eq. (1)) provides a CSR-normalised, scale-resolved measure of spatial interaction on networks, yielding a unified framework for unmarked, categorical, and continuously marked point patterns.

### Nonparametric bootstrap of correlation functions for spatial networks

To quantify uncertainty in netPCF estimates, we use a nonparametric spatial bootstrap framework that accounts for both finite sampling variation and the spatial dependence induced by the domain geometry. When spatial organisation is heterogeneous, empirical correlation estimates obtained from different regions of the domain may differ substantially. Spatial resampling therefore provides a practical means of quantifying the robustness of the estimated correlation structure.

We adopt Loh’s marked point bootstrap procedure for estimating uncertainty in correlation functions [35]. Briefly, the contribution of each node to the netPCF estimation, *g*^net^(*r*), is first computed on the initial data and stored. The spatial domain is then partitioned into subregions of equal volume and shape. Bootstrap estimates are generated by resampling these subregions with replacement, and computing *g*^net,∗^(*r*) from those residing within the sampled subregions. This process is repeated *m* times, generating a distribution of spatial bootstrap correlation estimates.

Confidence intervals are computed from the resulting bootstrap distribution using the basic bootstrap interval [20]. For a confidence level of 100(1 − *α*)%, this interval is given by

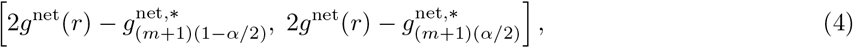

where *g*^net^(*r*) is the original estimate, 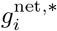, denotes the *i*th ordered bootstrap estimate.

A key requirement of this procedure is a partition of the spatial domain into contiguous regions of comparable volume and shape. While spatial partitioning methods are well established for Euclidean domains (domain tesselation), no standard approach exists for spatial networks. Therefore, we developed a node partition algorithm that divides the network into *k* subnetworks with approximately equal network volume (total edge length) while preserving spatial compactness. Full algorithmic details and validation are provided in Supplementary Information 1.4.

Using these partitions, we generate bootstrap replicates of the netPCF and thereby estimate spatial variability in the correlation function across the domain. Throughout this study, we report 95% (*α* = 0.05) confidence intervals obtained using this spatial bootstrap procedure for *m* = 1000.

### Comparing Pair Correlation Estimates

We quantify the similarity of pair correlation estimate curves using Normalised Cross-Correlation (NCC) [8]. The NCC is the Pearson correlation coefficient between the discretised pair correlation estimates evaluated over the distance range of interest, thereby quantifying similarity in their shape independently of magnitude. Subsequently, *NCC* ≈ 1 indicates strong agreement in correlation estimate curve shape, *NCC* ≈ 0 indicating no linear correspondence, and values *NCC* ≈ −1 indicating opposing trends between the two estimates.

## Results

### netPCF on Delaunay networks recapitulates classical PCFs in Euclidean space

To validate the network pair correlation function (netPCF), we first compared it against the extrinsic Euclidean-distance PCF using synthetic point processes with known second-order structure (see Supplementary Information 1.5 for details). A dense background point pattern generated from a homogeneous Poisson process was used to approximate the Euclidean domain, and a Delaunay network was constructed on the union of background and signal points to provide a distance-preserving discretisation of space. To avoid discretisation-induced bias in network estimates, kernel bandwidths were chosen to satisfy the sufficient-coverage criterion (Eq. (41)) derived in Supplementary Information 1.6.

Across all simulated processes, netPCF was in good agreement Euclidean PCF estimates (Fig. 4). This agreement was confirmed quantitatively using NCC, with all simulated examples producing high similarity between the netPCF and Euclidean PCF estimates (NCC *>* 0.88). For the homogeneous Poisson process, both estimators were consistent with complete spatial randomness, with *g*(*r*) ≈ 1 across distances (Fig. 4A). For inhibitory processes, including Matérn Type II and Strauss patterns [3], both methods recovered the expected short-range exclusion, characterised by *g*(*r*) *<* 1 at small distances before returning toward the null expectation at larger distances, and exhibiting periodicity, respectively (Fig. 4B–C). For the clustered Thomas process [3], both estimators detected elevated short-range correlation, with *g*(*r*) *>* 1 at small distances followed by decay toward *g*(*r*) ≈ 1 at larger distances (Fig. 4D).

**Fig 4.**
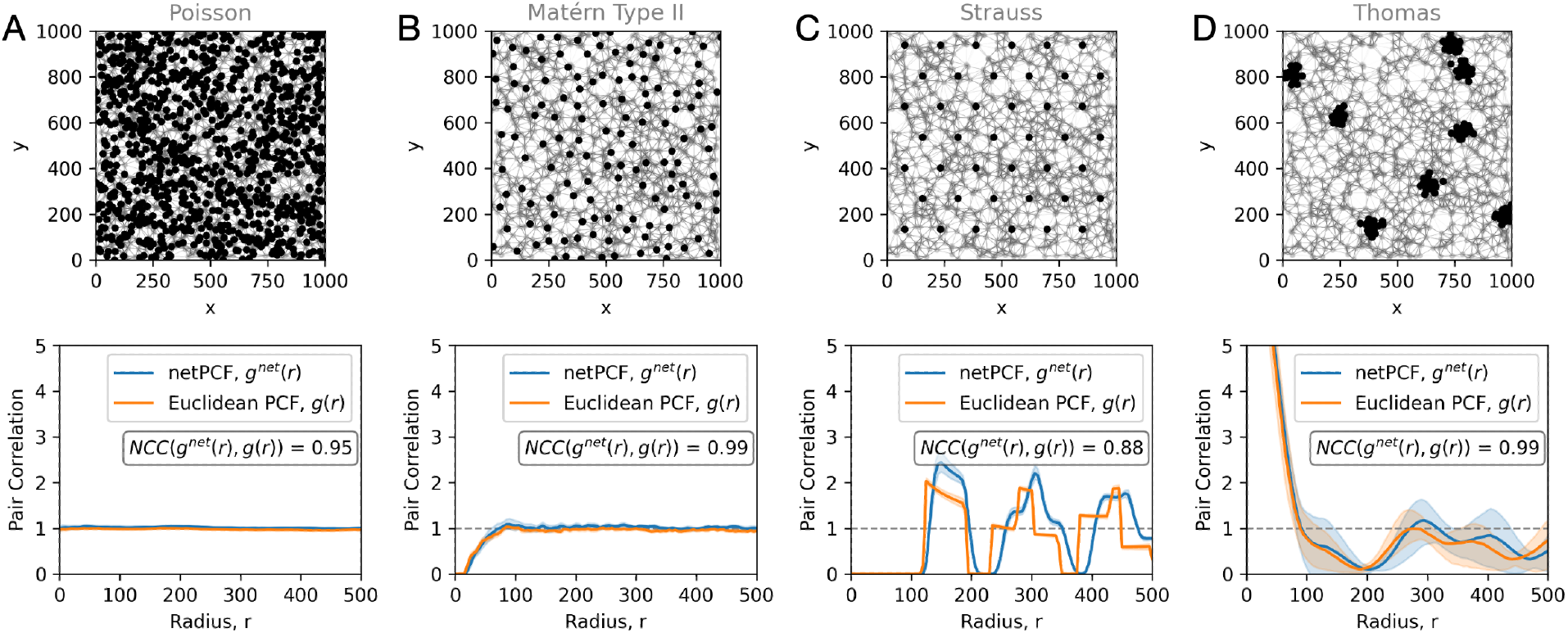
Recapitulating Euclidean PCF behaviour on spatial networks. Point processes embedded within a common spatial domain defined by a background Poisson point process (*ρ* = 0.001), from which an edge-filtered (*<* 100 *µ*m) Delaunay network is constructed using the union of all point patterns. (A) Poisson process (*ρ* = 0.001), (B) Matérn Type II process, (C) Strauss process, and (D) Thomas process. Corresponding Euclidean PCF and network-based PCF (netPCF) curves are shown beneath each network. Shaded regions correspond to 95% confidence interval about the estimated correlation. Normalised cross-correlation (NCC) values comparing Euclidean PCF and netPCF estimates are shown for each simulated process.

Together, these results show that sufficiently resolved Delaunay-based spatial networks preserve approximate extrinsic Euclidean distances for netPCF to reproduce classical PCF estimates in simple domains [48]. netPCF recovered the expected signatures of complete spatial randomness, short-range inhibition, and clustering, corresponding to spatial mixing, exclusion, and aggregation patterns commonly observed in biomedical imaging data [15, 41].

### Geometric constraints bias Euclidean PCFs but are resolved by netPCF

We next tested whether pair correlation estimates on spatial networks remain valid when when point patterns are constrained by the intrinsic geometry of curved or structured biological domains. This setting is common in spatial biomedical data, where cells may be confined by tissue boundaries [41], excluded from regions such as lumen [19], aligned along branching anatomical structures [52], or even distorted during image acquisition [32] (Fig. 1). In such cases, global Euclidean distance may misrepresent the spatial relationships available within the domain, causing geometry-induced effects to be mistaken for biological interaction.

To isolate this effect, we generated homogeneous Poisson point processes on three structured domains representing distinct geometric constraints: domains with excluded regions (*Holes*), branching domains (*Branching* ), and curved spiral (*Spiral* ) domains (Fig. 5A–C). Since points were sampled uniformly with respect to the intrinsic geometry of each domain, the expected pair correlation is complete spatial randomness, *g*(*r*) ≈ 1, across spatial scales. Deviations from this null therefore reflect bias introduced by the distance metric or domain representation, rather than true spatial interaction.

**Fig 5.**
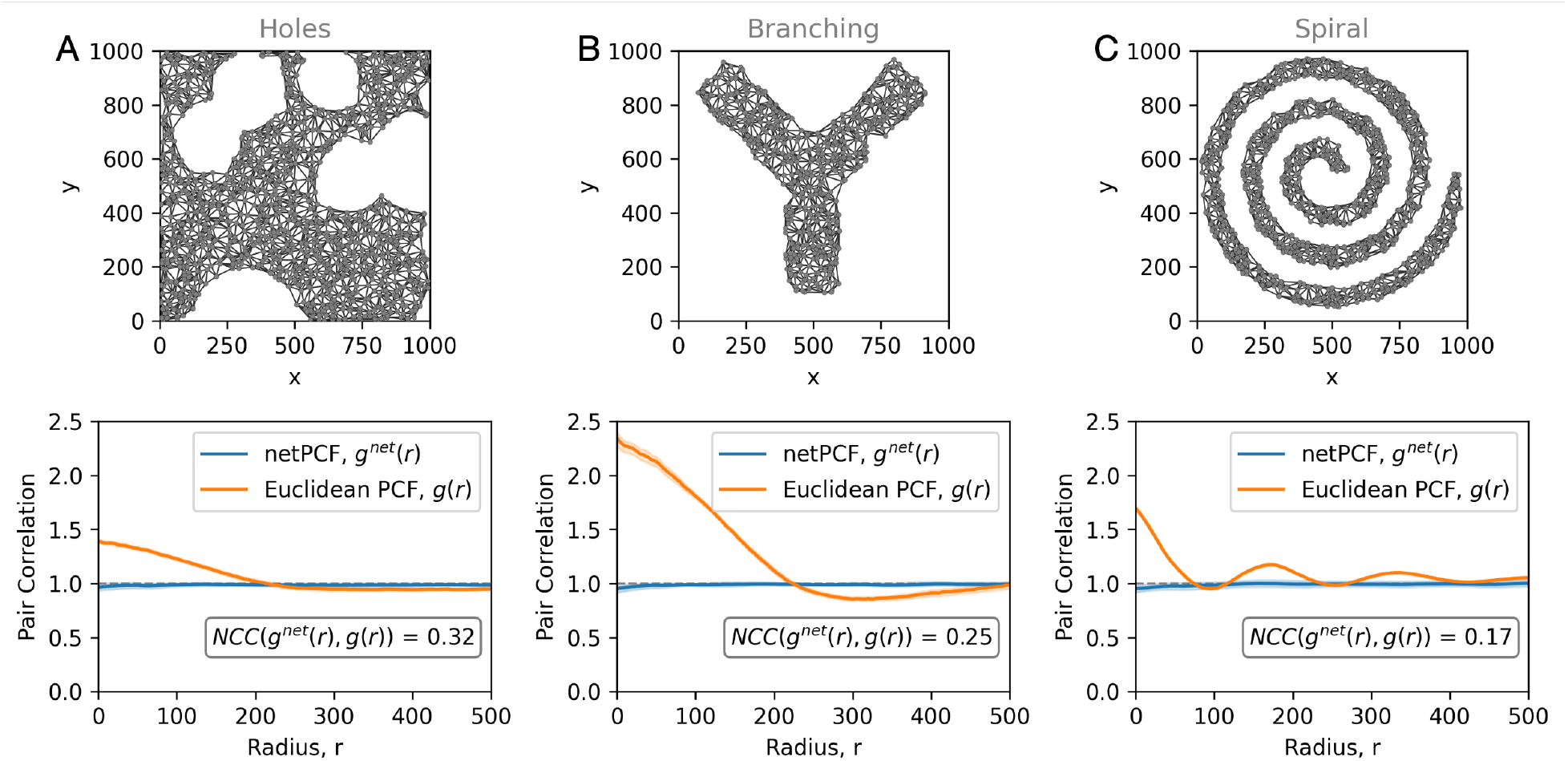
Intrinsic correction of domain geometry in spatial point patterns. Point patterns generated from a homogeneous Poisson process and embedded within structured domains that impose geometric constraints. (A) Domains with excluded regions (*Holes*), where large portions of space are inaccessible. (B) Branching domains (*Branching* ), where points are confined to a bifurcating structure. (C) Spiral domains (*Spiral* ), where points follow a curved trajectory. Corresponding Euclidean PCF and network-based PCF (netPCF) estimates for all nodes are shown beneath each domain. Shaded regions correspond to 95% confidence interval about the estimated correlation. Normalised cross-correlation (NCC) values comparing Euclidean PCF and netPCF estimates are shown for each domain.

Across all three examples, PCFs equipped with the extrinic Euclidean metric (Euclidean PCF) showed systematic departures from CSR even though the underlying process is random. The nature of the bias depended on the geometry. Excluded regions altered the available sampling space, branching structures imposed connectivity constraints, and curved domains made intrinsically distant regions appear artificially close in Euclidean space. Thus, Euclidean PCFs confounded geometric constraints with apparent clustering or inhibition.

In contrast, netPCF estimates remained consistent with CSR across all domains. By measuring approximate geodesic distances along the inferred spatial network, netPCF respects the intrinsic geometry of the domain and normalises by the accessible network length at each distance. Although Euclidean PCFs can incorporate accurate domain boundaries to account for finite support and edge effects, they remain defined with respect to global Euclidean distance. Consequently, even with accurate boundary information, they cannot capture the intrinsic separation of points constrained to curved or branching domains, where biologically relevant distances follow the domain rather than the embedding space. The resulting estimates differed substantially from the Euclidean PCF curves, with NCC values of 0.32, 0.25, and 0.17 for the *Holes, Branching*, and *Spiral* domains, respectively. These low NCC values indicate weak agreement between the Euclidean and network-based estimates, consistent with the Euclidean PCF detecting apparent spatial structure induced by domain geometry rather than by the point process itself. These results show that network-based PCFs can recover the correct null behaviour for point patterns constrained to perforated, branching, or curved domains, supporting the need for intrinsically geometry-aware spatial statistics in structured biological settings.

### Robustness of netPCF to Delaunay network construction and resolution

We next assessed how sensitive netPCF estimates are to uncertainty in Delaunay-based spatial network construction. In spatial imaging data, biological structures are captured, segmented, transformed into point clouds and, here, represented as spatial networks. Each step may introduce errors, sparse image resolution can reduce point density [2], segmentation or registration errors can perturb node positions [41, 10, 24], and network construction requires choices about the length scale of local connectivity [4].

To test whether these sources of uncertainty affect correlation estimates, we used a synthetic spiral domain with periodically placed nodes as a structured reference pattern. We then recomputed netPCF after perturbing the network construction in three ways: subsampling nodes, varying the maximum edge-length threshold, and adding spatial noise to node positions (Fig. 6A).

**Fig 6.**
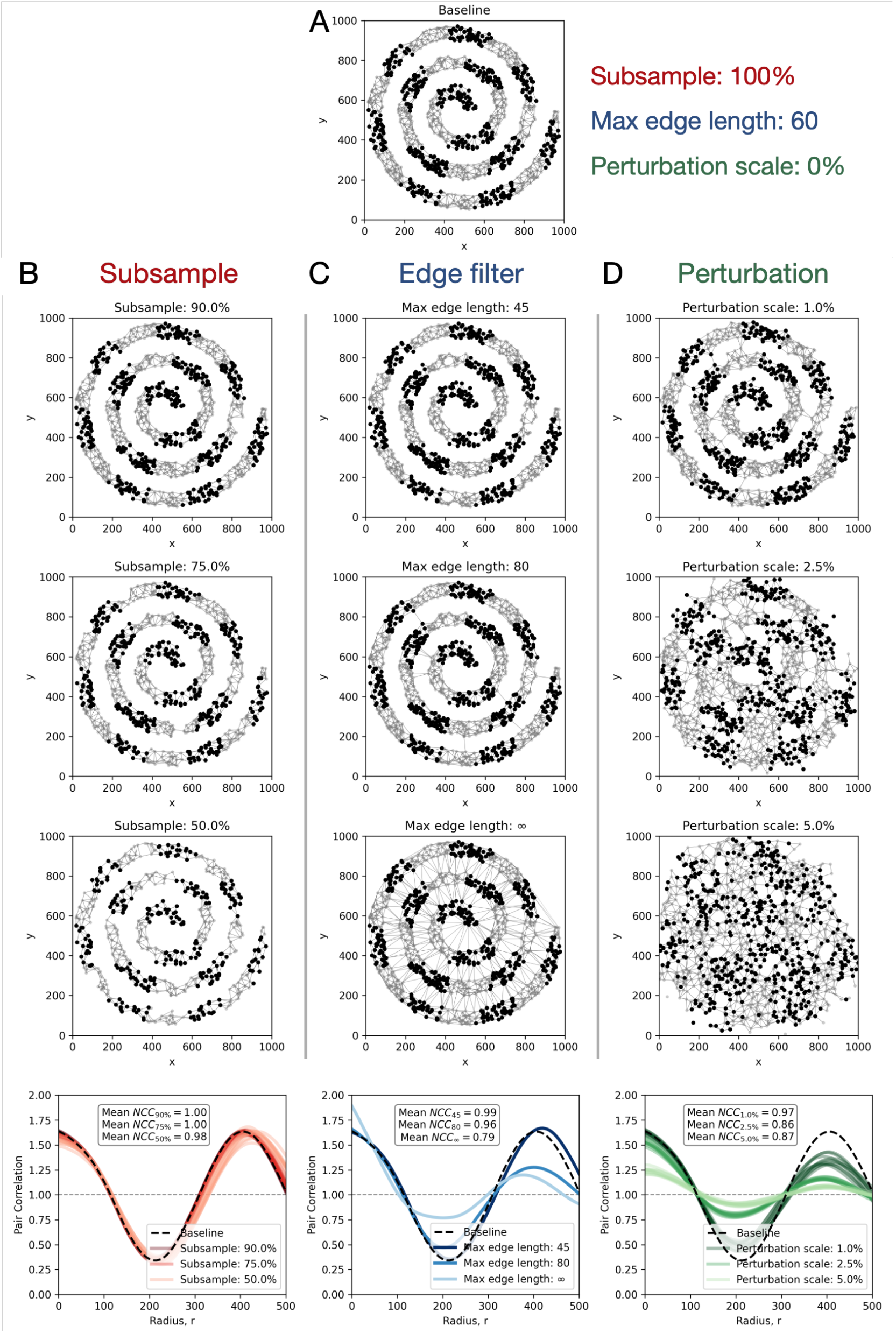
Robustness of netPCF to Delaunay-based domain estimation. (A) Baseline spiral domain constructed from a Poisson point process (*ρ* = 0.001), with periodic signal nodes shown in black. (B–D) Example re-estimated networks and corresponding netPCF curves after perturbing the baseline domain by subsampling points 90%, 75%, 50%, varying the maximum Delaunay edge-length threshold 45, 80, ∞, and adding Gaussian positional noise *σ* = 1%, 2.5%, 5% of domain width).Corresponding network-based PCF (netPCF) estimates for the black node populations are shown beneath each domain. Mean normalised cross-correlation (NCC) values compare perturbed netPCF estimates with the baseline across repeated trials.

First, we tested sensitivity to sampling density by randomly retaining 90%, 75%, or 50% of the baseline nodes. Across repeated subsamples, netPCF estimates remained highly similar to the baseline estimate (Fig. 6B), with mean NCC values exceeding 0.98 across all subsampling levels. This indicates that netPCF is robust to moderate reductions in node density, provided that sampling preserves the connectedness of the network.

Second, we varied the maximum edge length used to filter the Delaunay network. Estimates from the baseline network were highly consistent with those obtained using the shortest threshold that preserved network connectivity (Fig. 6C). Increasing the threshold, or retaining the full unfiltered Delaunay triangulation, introduced longer-range edges that increasingly shortcut the intrinsic spiral geometry. This reduced the fidelity of the network representation and dampened the amplitude of the netPCF signal. Nevertheless, the qualitative periodic structure was preserved across all thresholds, with NCC values exceeding 0.79 relative to the baseline estimate.

Finally, we examined sensitivity to positional noise by perturbing node locations with Gaussian noise of increasing magnitude. Small perturbations produced netPCF estimates that closely matched the baseline, whereas larger perturbations attenuated longer-range periodic structure (Fig. 6D). Short-range correlations remained comparatively stable, while long-range features degraded as spatial perturbations increasingly distorted the inferred domain geometry. Despite this attenuation, the qualitative periodic structure was preserved across all noise levels, with NCC values exceeding 0.87 relative to the baseline estimate.

Collectively, these analyses show that netPCF is robust to moderate variation in Delaunay network construction, including reduced sampling density, reasonable changes in edge-length filtering, and positional noise (further quantitative comparisons in Supplementary Information 1.8). Deviations become substantial only when the network no longer preserves the intrinsic geometry of the domain. This supports the use of Delaunay-based spatial networks as a stable representation for geometry-aware pair correlation analysis.

### Categorical and continuous extensions of netPCF in arbitrary spatial dimensions

Spatial biomedical datasets contain information more than object locations: nodes may also carry categorical attributes, such as cell type or tissue compartment, and continuous attributes, such as protein intensity, transcript abundance, or morphological measurements [15]. In these settings, the aim is often not only to quantify the spatial structure of the point pattern itself, but to measure how node-level features are organised over the underlying tissue geometry. We therefore extend netPCF from unmarked point patterns to marked spatial data, allowing spatial correlations to be estimated for both categorical and continuous node attributes [13, 3]. Rather than requiring separate estimators for different mark types or spatial geometries, the same network-based formulation is recovered by changing only the feature kernel in Eq. (1). Full definitions of the corresponding feature kernels are provided in Supplementary Information 1.3.

We first considered categorical marks using the cross-netPCF, where feature kernels are indicator functions selecting pairs of node classes. Across three synthetic domains of increasing geometric complexity, a two-dimensional spiral (Fig. 7A), a cylindrical surface embedded in three dimensions (Fig. 7B), and a three-dimensional sphere (Fig. 7C), cross-netPCF recovered the imposed periodic association between categorical labels. In each case, the estimator detected alternating spatial co-localisation and exclusion between node types, while respecting the intrinsic geometry of the domain as exhibited by netPCF estimation of all nodes.

**Fig 7.**
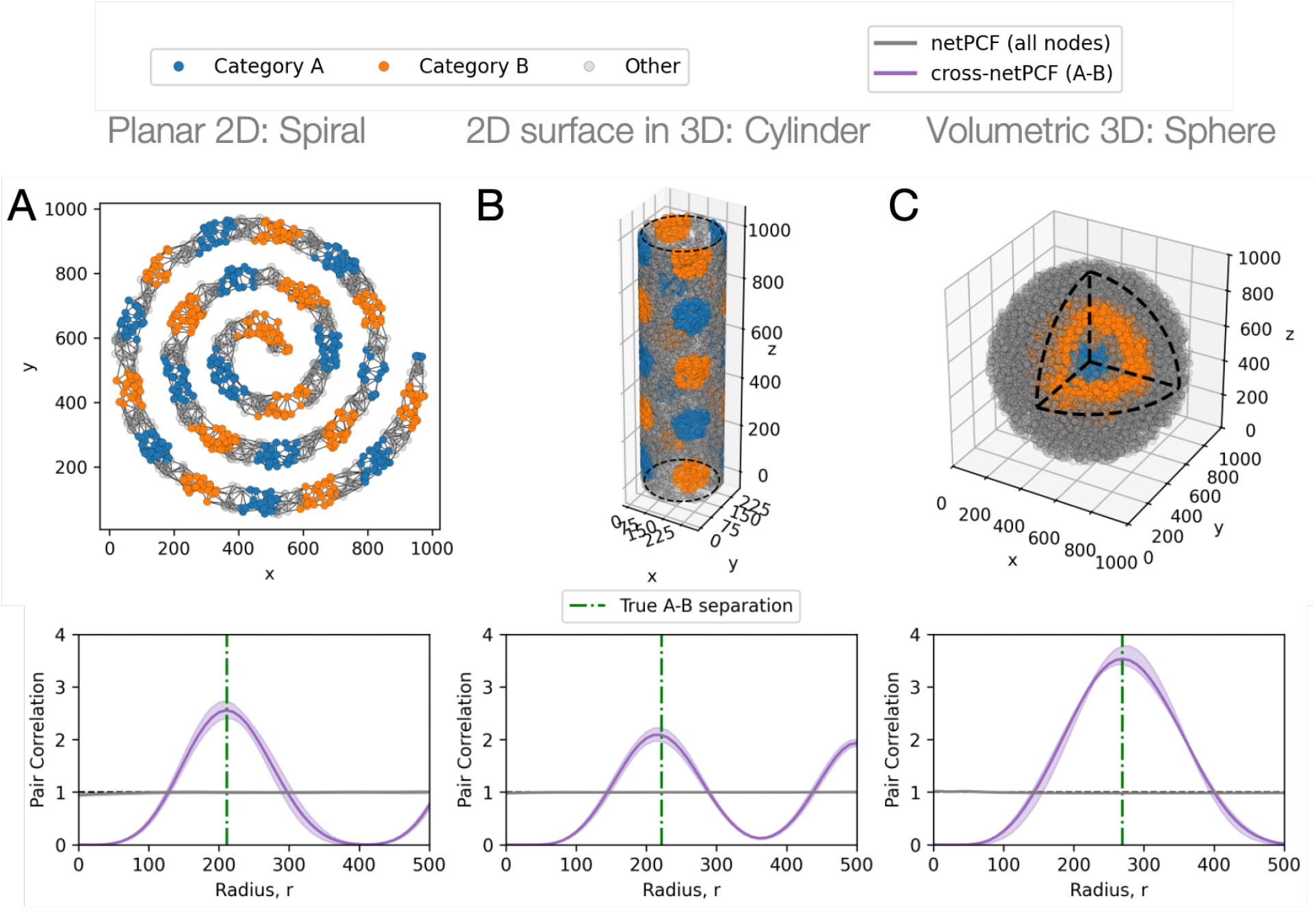
Quantifying categorical data on spatial networks. Synthetic point patterns with nodes assigned categorical labels (A or B) across domains of increasing geometric complexity. (A) Two-dimensional spiral domain, where points are generated from a Poisson point process along a curved trajectory and labels A and B are assigned periodically along the spiral. (B) Two-dimensional surface embedded in three dimensions, defined by a hollow cylindrical domain (radius 137, height 1000), with points sampled from a Poisson point process over the surface and labels assigned periodically. (C) Three-dimensional volumetric domain, where points are generated from a Poisson point process within a sphere (radius 500) and labels are assigned periodically as a function of radial distance from the centre. Cross network-based PCF (netPCF) estimates between nodes of type A and B are shown in purple beneath each domain, while netPCF estimates for all nodes are shown in grey. Shaded regions denote 95% confidence intervals about the estimated correlations.

We then considered continuous node attributes using the weighted-netPCF, where feature kernels weight nodes according to similarity to a target continuous value (see Supplementary Information 1.3) [11, 13]. This continuous extension similarly recovered imposed periodic mark structure across the same classes of domains, including curved, surface, and volumetric geometries (Supplementary Information 1.7). Together with the categorical case, this shows that netPCF provides a unified estimator for unmarked, categorical, and continuous marked point processes in arbitrary spatial dimensions.

### Distinguishing geometry-driven and intrinsic endothelial–immune interactions in the 3D tumour microenvironment

Spatial relationships between endothelial and immune cells are central to tumour biology because they may reflect immune infiltration, vascular access for extravasation and metastasis, and the organisation of immune niches within the tumour microenvironment. In HER2+ breast carcinoma, where immune composition and spatial organisation can inform tumour state and therapeutic response [5], distinguishing genuine immune–vascular organisation from tissue-architecture effects is therefore important. However, the geometry of the sampled domain can generate apparent cellular proximity in structured tissues. Previous analysis of 3D imaging mass cytometry (IMC) data showed that endothelial–immune distances differ between serial 2D sections and full volumetric reconstructions, demonstrating that dimensionality and tissue geometry affect spatial inference [31].

We re-analysed this previously published HER2+ breast carcinoma IMC dataset, which contains endothelial cells and immune subsets including B cells, CD8+ T cells, and CD8 − T cells (Fig. 8A) [31]. From nuclear segmentation and phenotypic classification, we obtained single-cell 3D point clouds describing the spatial organisation of these populations within the reconstructed tissue volume (Fig. 8B). We first reproduced the reported difference in nearest-neighbour endothelial–immune distances between serial 2D sections and the full 3D reconstruction (Fig. 8C). We then used netPCF to extend this analysis from proximity alone to scale-dependent spatial association, asking whether endothelial–immune organisation is preserved after accounting for the geometry of the tumour microenvironment.

**Fig 8.**
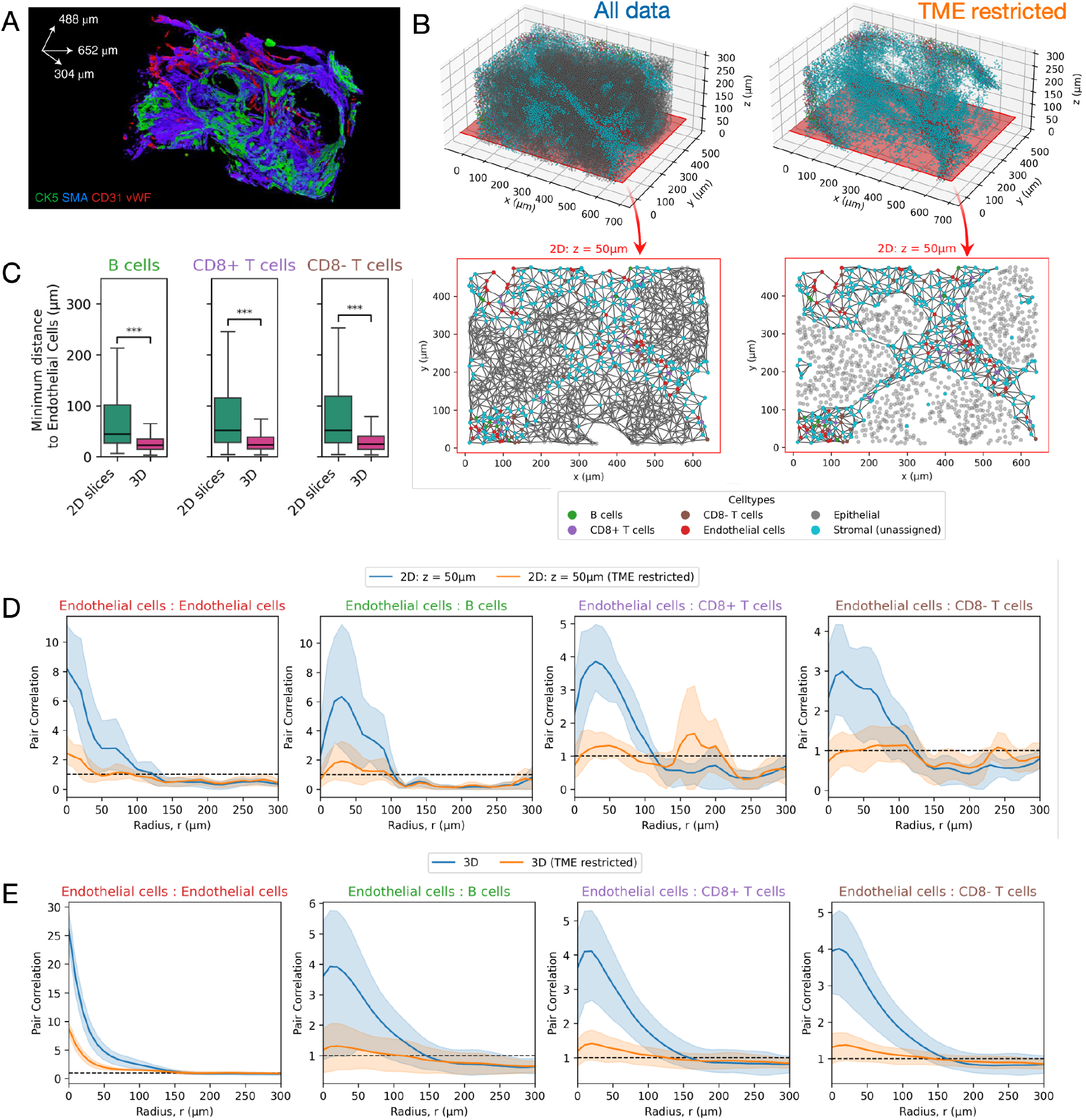
Geometry-aware quantification of endothelial–immune interactions in the breast cancer tumour microenvironment. (A) Three-dimensional rendering of imaging mass cytometry data from a breast carcinoma sample, showing expression of basal markers (CK5, SMA) and endothelial markers (vWF, CD31). (B) Reconstructed 3D point cloud following segmentation and cell type classification, shown for the full dataset (All data) and with epithelial cells removed (TME restricted). A representative 2D section (*z* = 50–54 *µ*m) is shown for both cases, together with the corresponding spatial networks defining the domains of analysis. (C) Distribution of minimum distances from immune cell subsets to endothelial cells, comparing measurements across serial 2D sections (4 *µ*m spacing) and the full 3D dataset. Statistical significance is assessed using the Mann–Whitney U test (^∗∗∗^*p <* 0.001). (D) Network-based cross pair correlation functions (netPCF) between endothelial cells and immune cell subsets (B cells, CD8+ T cells, and CD8-T cells) computed on a representative 2D section (*z* = 50 *µ*m), shown for both full and TME-restricted domains. (E) Corresponding netPCF estimates computed on the full 3D point cloud, for both full and TME-restricted domains. Shaded regions in (D) and (E) denote 95% confidence intervals.

To separate these effects, we compared endothelial–immune associations across two spatial domains. The first was the full tissue domain, containing all segmented cells. The second was a TME-restricted domain, constructed after removing epithelial tumour cells. This restriction conditions the analysis on the non-epithelial tumour microenvironment and tests whether apparent endothelial–immune associations persist once large-scale tumour architecture is removed from the spatial support. We computed cross-netPCFs between endothelial cells and each immune subset in both domains, using both a representative 2D section and the full 3D reconstruction (Fig. 8D–E).

Endothelial–endothelial netPCFs showed robust short-range clustering in both the full and TME-restricted domains, consistent with preservation of vascular structure under domain restriction. Endothelial–immune associations were more domain-dependent. In the representative 2D section, endothelial–immune netPCFs showed apparent short-range co-localisation in the full domain, with partial attenuation after TME restriction (Fig. 8D). Some qualitative structure persisted between domains, including longer-range endothelial–B cell exclusion, indicating that 2D analysis alone did not fully resolve the contribution of tissue geometry to the observed associations.

The distinction became clearer in 3D. In the full 3D tissue domain, endothelial cells appeared positively associated with immune subsets at short range. After restricting the analysis to the TME domain, this apparent endothelial–immune co-localisation was removed, and endothelial–immune netPCFs were consistent with spatial randomness across scales (Fig. 8E). Thus, in this HER2+ breast carcinoma sample, apparent immune–vascular proximity is largely explained by the global architecture of the tumour and surrounding tissue compartments, rather than by a strong intrinsic endothelial–immune spatial association within the TME. This interpretation is consistent with the hypervascular and spatially heterogeneous stroma often associated with HER2+ breast carcinoma, where pro-angiogenic signalling can generate endothelial-dense regions that structure the local tissue environment [44]. The observed endothelial–immune proximity therefore appears more consistent with shared localisation within stromal architecture than with a robust local immune–vascular interaction.

Importantly, this was not due to a general loss of spatial signal after domain restriction. Immune–immune associations showed a different pattern: short-range co-localisation between immune subsets was preserved within the TME-restricted domain, including B cell–CD8+ T cell co-localisation (Supplementary Information 1.9). This suggests that, once tumour geometry is accounted for, the dominant local immune structure in this sample is not immune proximity to vasculature, but immune aggregation within the TME. In HER2+ breast carcinoma, the spatial organisation of B cells and CD8+ T cells has been associated with favourable clinical outcome and anti-tumour immune activity [61]. More recent work has highlighted B cell spatial organisation, including their relationship to invasive tumour regions and endothelial compartments, as a potential marker of immune activation and immunotherapy-relevant CD8+ T cell function [5, 43]. The persistence of B cell–CD8+ T cell co-localisation after TME restriction therefore supports the presence of a locally organised immune compartment in this sample. By contrast, the weak endothelial–immune signal after accounting for tissue geometry suggests limited evidence for spatially localised immune–vascular coupling, indicating that immune aggregation within the TME is the more robust spatial feature.

These analyses show how geometry-aware pair correlation can refine the biological interpretation of spatial organisation in complex 3D tissues. Conventional nearest-neighbour summaries identify close cell pairs, but they do not determine whether proximity reflects local cellular organisation or the shape of the sampled tissue. By combining scale-resolved correlation estimates with explicit spatial domain restriction, netPCF distinguishes associations that are preserved within the TME from those induced by tissue architecture. In this HER2+ breast carcinoma sample, this separates geometry-driven endothelial–immune proximity from intrinsic immune aggregation, providing a more reliable framework for interpreting spatial associations in 3D tumour biology.

### Quantifying multiscale spatio-temporal gene pattern transitions in a developing mammalian embryo

We next applied netPCF to Wnt gene expression patterns across the surface of developing murine embryos. Wnt genes regulate embryonic patterning, tissue specification, and organogenesis [34], and their expression domains are organised over curved, evolving surfaces where Euclidean distance may poorly capture intrinsic spatial relationships (Fig. 1E). We therefore re-analysed the 3D spatiotemporal Wnt signalling atlas of Murphy et al. [42], which mapped Wnt gene expression to embryonic spatial reference models at E9.5 and E11.5. These stages capture a developmental transition from broad expression domains during early organogenesis to more intricate boundaries and regions of high Wnt occupancy later in development [34, 42]. Building on the original voxel-based co-expression spatial analysis [42], we used netPCF to quantify multiscale spatial associations in Wnt expression across the embryonic surface during development.

Embryonic surface networks were reconstructed from processed Optical Projection Tomography (OPT) surface masks at E9.5 and E11.5, with each node assigned both continuous Wnt expression marks and binary expression labels (see Supplementary Information 1.10). This enabled netPCF analysis of both categorical and continuous expression of *Wnt1, Wnt2*, and *Wnt6* (Fig. 9A), genes associated with epithelial and surface-linked (ectodermal) developmental patterning [16].

**Fig 9.**
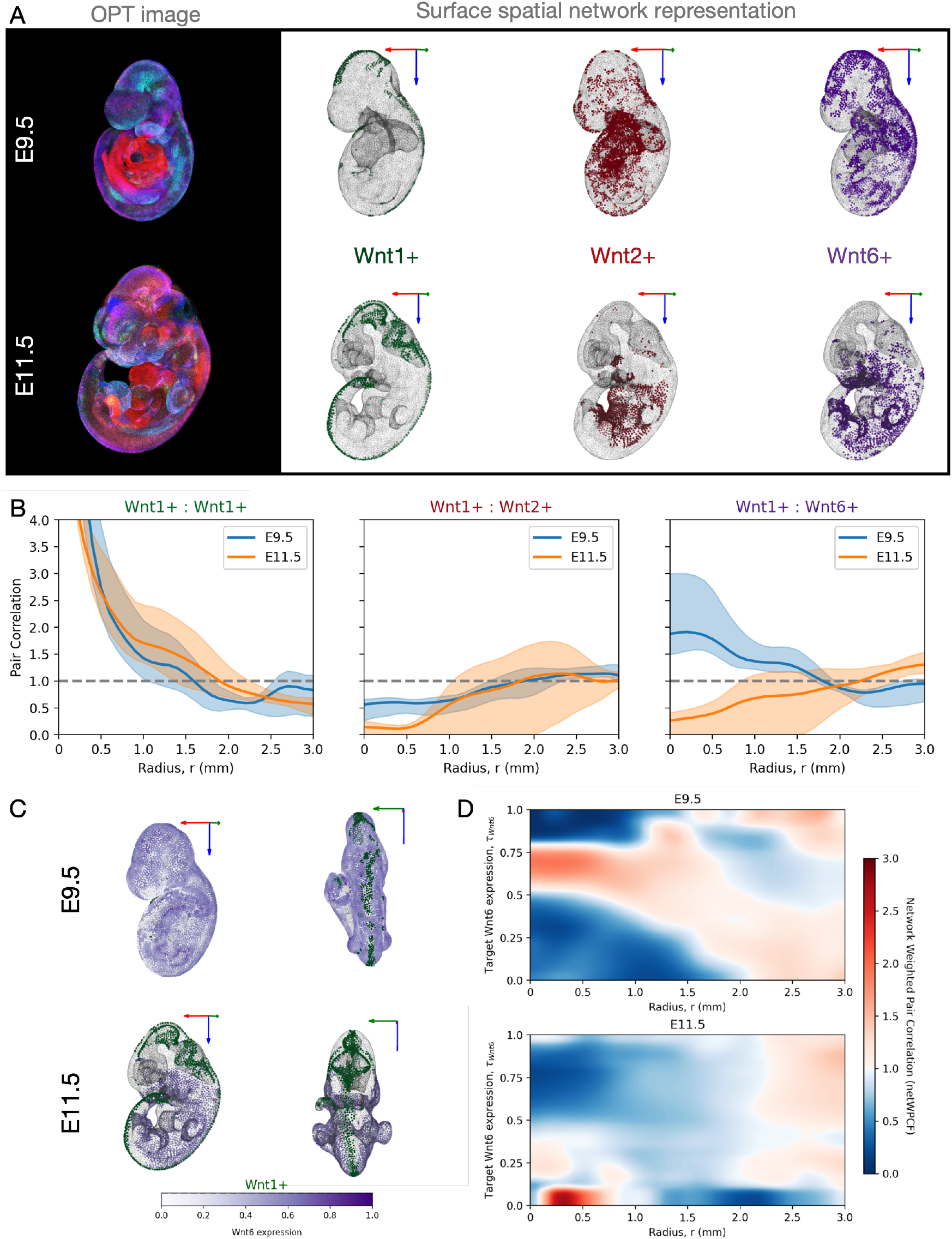
Spatio-temporal analysis of Wnt expression patterns on the surface of a developing murine embryo. (A) Spatial gene expression patterns from the Wnt and Fzd families in E9.5 and E11.5 murine embryos imaged by optical projection tomography. Embryo surface networks are shown with nodes highlighted according to categorical activation of selected Wnt genes. Red, green, and blue arrows indicate the orientation and scale of the x-, y-, and z-axes, respectively, with each arrow corresponding to 1 mm. (B) Cross-netPCF curves between *Wnt1, Wnt2*, and *Wnt6* expression domains computed over the surface networks of E9.5 and E11.5 embryos. Shaded regions denote 95% confidence intervals. (C) Surface networks of the E9.5 and E11.5 embryos coloured by categorical *Wnt1* expression and continuous *Wnt6* expression. Red, green, and blue arrows indicate the orientation and scale of the x-, y-, and z-axes, respectively, with each arrow corresponding to 1 mm. (D) Weighted netPCF heatmaps between *Wnt1* + nodes and continuous *Wnt6* surface expression for E9.5 and E11.5 embryos, evaluated across all target *Wnt6* expression values from to 1. Blue regions indicate spatial exclusion, whereas red regions indicate co-localisation.

We first computed cross-netPCF estimates between categorical expression domains for *Wnt1, Wnt2*, and *Wnt6* at both E9.5 and E11.5 (Fig. 9B). Across developmental stages, *Wnt1* showed consistent scale-dependent relationships with itself and with *Wnt2*. Namely, *Wnt1* –*Wnt1* estimates exhibited short-range co-localisation, whereas *Wnt1* –*Wnt2* estimates showed spatial exclusion. These patterns are consistent with the interpretation that *Wnt1* occupies a spatially restricted expression domain that remains distinct from the ventral territories associated with *Wnt2* as demonstrated by Murphy et al. [42].

In contrast, the spatial relationship between *Wnt1* and *Wnt6* appeared to change markedly between developmental stages. At E9.5, cross-netPCF estimates indicated short-range enrichment between *Wnt1* - and *Wnt6* -positive regions, consistent with these expression domains being spatially adjacent, or locally interspersed, on the embryonic surface. By E11.5, this relationship shifted towards short-range exclusion, suggesting increased spatial segregation between the two domains as development proceeds. In prior analysis, no voxel-wise co-localisation or exclusion of *Wnt6* with other Wnt genes, including *Wnt1*, was identified at any developmental stage [42].

To extend this analysis beyond categorical expression states, we next applied the weighted-netPCF (netWPCF), using *Wnt1* -positive nodes as the reference population and continuous normalised *Wnt6* expression as the target mark (Fig. 9C–D). At E9.5, *Wnt1* -positive nodes were short-range co-localised with nodes exhibiting high *Wnt6* expression, approximately 0.7 in normalised expression units, over distances below 0.75 mm. These same *Wnt1* -positive nodes were excluded from regions with lower *Wnt6* expression. Moreover, the associated *Wnt6* expression level decreased with increasing surface distance from *Wnt1* -positive nodes, indicating a decay in *Wnt6* activity with distance from the *Wnt1* domain. By contrast, at E11.5, *Wnt1* -positive nodes were preferentially associated only with regions of low or near-zero *Wnt6* expression, below approximately 0.2 normalised expression units, while being excluded from regions with higher *Wnt6* expression. This continuous-mark analysis supports the categorical netXPCF result, suggesting a developmental transition from *Wnt1* –*Wnt6* co-localisation to exclusion, robust to expression thresholding.

These observations are consistent with the classic model of ectodermal boundary refinement during neural tube closure between E9.5 and E10.5, where initial *Wnt1* -*Wnt6* overlap induces neural crest specification before long-range inhibitory signaling prompts a medial-to-lateral clearance of *Wnt6*, thereby segregating the *Wnt1* -positive dorsal neuroectoderm from the overlying non-neural surface ectoderm [26, 23]. Critically, by resolving Wnt expression patterns over the intrinsic geometry of the embryonic surface, netPCF reveals surface expression dynamics not detected by voxel-based methods and recovers established relationships between Wnt genes in the developing ectoderm.

## Discussion

We introduced netPCF, a network-based extension of the pair correlation function for point patterns defined on data-driven spatial domains. By substituting the extrinsic Euclidean distance for approximated geodesic distance on a reconstructed spatial network, netPCF extends classical spatial statistics to settings in which the geometry of the underlying domain is irregular, curved, or only implicitly observed. This provides a principled way to quantify spatial organisation in biological data where the relevant spatial substrate is not well described by a simple planar metric. In doing so, netPCF addresses an important gap between the rich geometric complexity of modern spatial omics and imaging data, and the tools available for analysing spatial dependence on such domains.

A central advantage of netPCF is that it helps distinguish biological spatial interaction from geometry-induced proximity in complex spatial data. In irregular domains, apparent clustering or inhibition may arise not only from biological organisation, but also from constraints imposed by the shape, curvature, or connectivity of the tissue itself. By representing the domain as a spatial network and defining correlation relative to network geodesics, netPCF mitigates this source of bias and enables multiscale spatial structure to be interpreted in a geometry-aware manner. This was illustrated in the breast carcinoma example, where spatial relationships in the tumour microenvironment were strongly conditioned by tissue architecture. In the developmental context, netPCF further revealed previously overlooked spatial correlations between *Wnt1* and *Wnt6* by accounting for the intrinsic geometry of the embryonic surface. These examples highlight the importance of representing spatial context accurately when interpreting biological patterning, and demonstrate how netPCF provides a principled and practical framework for estimating pair correlations in complex spatial biological domains.

Methodologically, netPCF extends the literature on spatial statistics on networks beyond the classical setting of fixed linear networks. Prior work has largely focused on one-dimensional network structures such as roads, river systems, or fibre-like domains [4, 3, 1]. In contrast, netPCF is designed for arbitrary spatial graphs reconstructed from data, making it suitable for biological systems in which the domain is not known a priori and must itself be inferred. This generality is important for applications in spatial biology, where tissue surfaces, volumetric structures, and complex interfaces often need to be represented by meshes, graphs, or other discrete approximations before spatial statistics can be meaningfully applied.

The framework is also flexible across data types and geometric settings. Although we have emphasised point patterns, the same representation can accommodate planar domains, curved surfaces, and volumetric structures, as well as categorical or continuous node attributes. This makes netPCF broadly applicable to spatial-omics and other emerging modalities that generate highly resolved cellular and subcellular data across large tissue regions, where spatial organisation is central to interpretation [33]. More broadly, the method provides a unified language for geometry-aware summary statistics on irregular domains, and may help bridge the methodological divide between spatial statistics, computational geometry, and spatial systems biology.

However, netPCF depends on an accurate reconstruction of the domain geometry. As with any method based on an inferred representation, errors in network estimation may propagate into the geodesic distance metric and therefore into the estimated correlation function. We found the approach to be robust to sampling density, edge filtering, and noise in Delaunay-based constructions, but the choice of network parameters remains important. In many applications, the true domain is not directly known, which limits the extent to which these choices can be tuned a priori. This motivates future work on automatic domain detection, including machine learning approaches to manifold estimation and mesh reconstruction, as well as the use of alternative planar or surface-based network constructions. The embryo example presented, used nodes derived from an exterior contour mesh, illustrating how such flexibility may be useful in practice, but it also underscores the need for careful domain-specific validation.

Another important limitation is the choice of null model. Here we used complete spatial randomness as a practical baseline, because it provides a clear and interpretable reference point for identifying departure from spatial neutrality. From the perspective of spatial point process theory, however, the netPCF is a second-order summary statistic that is applicable to arbitrary point processes defined on the underlying manifold or network; the choice of CSR simply specifies the reference model against which departures are assessed. Many biological tissues are spatially inhomogeneous by construction, and point patterns on prescribed networks are often similarly nonstationary [13, 1]. In such cases, CSR may be overly simplistic, and model-based or inhomogeneous null processes will be needed to separate genuine interaction from large-scale intensity variation [18, 39]. Extending netPCF to these settings is an important direction for future work, particularly for data in which spatial heterogeneity is itself a dominant biological signal.

From a computational perspective, the current implementation of netPCF in `spacenet` is tractable for a wide range of imaging datasets, including modern spatial transcriptomics and proteomics applications [9]. We have taken care to support practical use on standard computing hardware through careful implementation, unit testing, and documentation. Nevertheless, further gains in efficiency and scalability are possible. In particular, accelerated kernel evaluation methods, including FFT-based approaches [50], and lower-level high-performance implementations, could support larger datasets and more demanding analyses. These developments would be especially valuable as spatial technologies continue to scale in both resolution and sample size.

The network formulation also creates a natural foundation for extending beyond pair correlation functions. In particular, analogous network-based versions of K-functions [47], mark correlation functions [39], and partition-based summaries could provide a broader suite of geometry-aware spatial statistics. In particular, our partition framework facilitates regional multi-species analyses used within spatial biology [15], including quadrat correlation analysis [40] and the Morisita-Horn index [28] on spatial networks. More generally, these extensions suggest the possibility of a unified statistical toolkit for analysing spatial structure across both continuous and discrete representations of geometry.

Overall, netPCF represents a step towards a more general framework for spatial analysis on irregular, inferred, and data-driven domains. As spatial technologies increasingly generate high-resolution measurements in geometrically complex settings, methods that explicitly account for domain structure will become increasingly important. By incorporating geometry into the definition of spatial dependence, netPCF provides a flexible and statistically grounded approach for studying multiscale organisation in spatial biology.

## Conclusion

In conclusion, netPCF extends the classical pair correlation function to spatial data defined on complex, data-driven domains by incorporating spatially-resolved network geodesics into the estimation of spatial dependence. This geometry-aware formulation allows intrinsic spatial organisation to be distinguished from artefacts of domain shape, making it particularly well suited to modern spatial biology datasets in which tissue architecture, curvature, and irregular boundaries are fundamental to interpretation. By providing a statistically grounded framework for analysing marked (labelled) point patterns on reconstructed spatial networks, netPCF offers a flexible foundation for multiscale spatial analysis independent of spatial dimension, across planar, surface, and volumetric settings. More broadly, it contributes toward a unified approach to geometry-aware spatial statistics on irregular and inferred domains, with clear opportunities for extension to additional summary functions, alternative null models, and learned spatial representations of data.

## Code Information and Availability

All methods introduced in this study are implemented in the open-source Python package `spacenet` (v0.0.2). The package provides a framework for constructing and analysing spatial networks. Comprehensive documentation for `spacenet`, including installation instructions, API reference, and usage examples, is available at https://www.spacenet-python.com. netPCF and the extended forms are implemented within the `spacenet.point patterns` submodule. The repository for `spacenet` can be found at https://github.com/joshwillmoore1/spacenet.

A repository that includes reproducible scripts and demonstration notebooks that replicate the synthetic and biological experiments presented in this study can be found at https://github.com/joshwillmoore1/netPCF-analysis.

The core functionality developed here is integrated into the `MuSpAn` Python package (version vX.X.X onward), which provides a broader framework for spatial data analysis. The `MuSpAn` platform includes additional tutorials and workflows demonstrating netPCF application.

## Data Information and Availability

All synthetic and processed data used in this study can be found at https://github.com/joshwillmoore1/netPCF-analysis. Information and source data for the IMC breast carcinoma data can be found at [31]. Information and source data for the murine embyro dataset can be found at [42].

## Acknowledgments

JWM and JAB were supported by Cancer Research UK (CRUK) grant number CTRQQR-2021 \ 100002, through the Cancer Research UK Oxford Centre.

## Author Contributions

JWM: Conceptualization, Methodology, Software, Investigation, Data Curation, Writing – Original Draft. JAB: Writing – Review & Editing. HMB: Writing – Review & Editing. All authors reviewed, approved, and agreed to the published version of the manuscript.

## 1 Supplementary Information

### 1.1 Polynomial kernels with finite support

In this section, we provide additional justification for the choice of spatial kernel used in Eq. (3), and discuss the properties of the family of polynomial kernels employed in this study.

We consider a class of compactly supported polynomial kernels *κ*_Δ*r,n*_(*x*) (Eq. (3)), and redefined here,

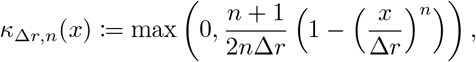

parameterised by bandwidth Δ*r* and polynomial degree *n* ∈ ℝ_*>*0_. These kernels provide a flexible mechanism for weighting pairwise distances, allowing interactions to be localised within a neighbourhood of radius Δ*r* while maintaining smooth behaviour within this region.

By construction, *κ*_Δ*r,n*_(*x*) has finite support for |*x*| *<* Δ*r*. This locality is particularly important for computational efficiency and interpretability, as it restricts contributions to pairs of nodes within a bounded distance, and ensures that estimates reflect interactions within a well-defined spatial scale.

The kernel is normalised such that

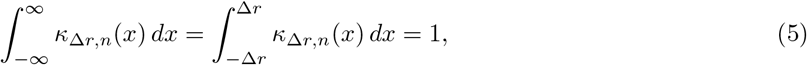

ensuring that it defines a valid weighting function over distances and yields consistent estimates under the complete spatial randomness (CSR) null model.

The degree parameter *n* controls the shape and smoothness of the kernel. For *n* = 1, the kernel is linear (Triangle kernel), while *n* = 2 is parabolic (Epanechnikov kernel), a widely used choice in kernel density estimation and spatial statistics as it minimises the Mean Integrated Squared Error [21, 3]. Increasing *n* leads to progressively flatter kernels, and in the limit as *n* → ∞, the kernel converges to a uniform (top-hat) function,

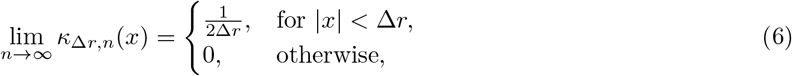

which corresponds to uniform weighting within an annulus of width 2Δ*r*.

This family of kernels provides a continuous interpolation between sharply localised weighting and uniform binning, controlled by the exponent *n*. For *n* ∈ (0, 1), the kernel mass is concentrated near the target distance *r*, placing greater emphasis on pairwise contributions close to this value. As *n* increases (*n >* 1), the kernel becomes progressively flatter within its support, distributing weight more uniformly across the bandwidth and reducing sensitivity to small deviations in distance. In the limit of large *n*, the kernel approaches a uniform (top-hat) weighting.

This flexibility allows control over the bias–variance trade-off in the estimation of pair correlation functions. In practice, moderate values of the exponent, *n* ∈ [0.2, 5], provide a balance between smoothness and localisation, while retaining computational efficiency due to finite support.

In this study, we use *n* = 10 when aiming to approximate uniform kernel behaviour, consistent with standard implementations of the Euclidean PCF ([15]), and *n* = 2 otherwise, corresponding to the Epanechnikov kernel. Examples of these kernels for different values of *n* are shown in Fig. 10.

**Fig 10.**
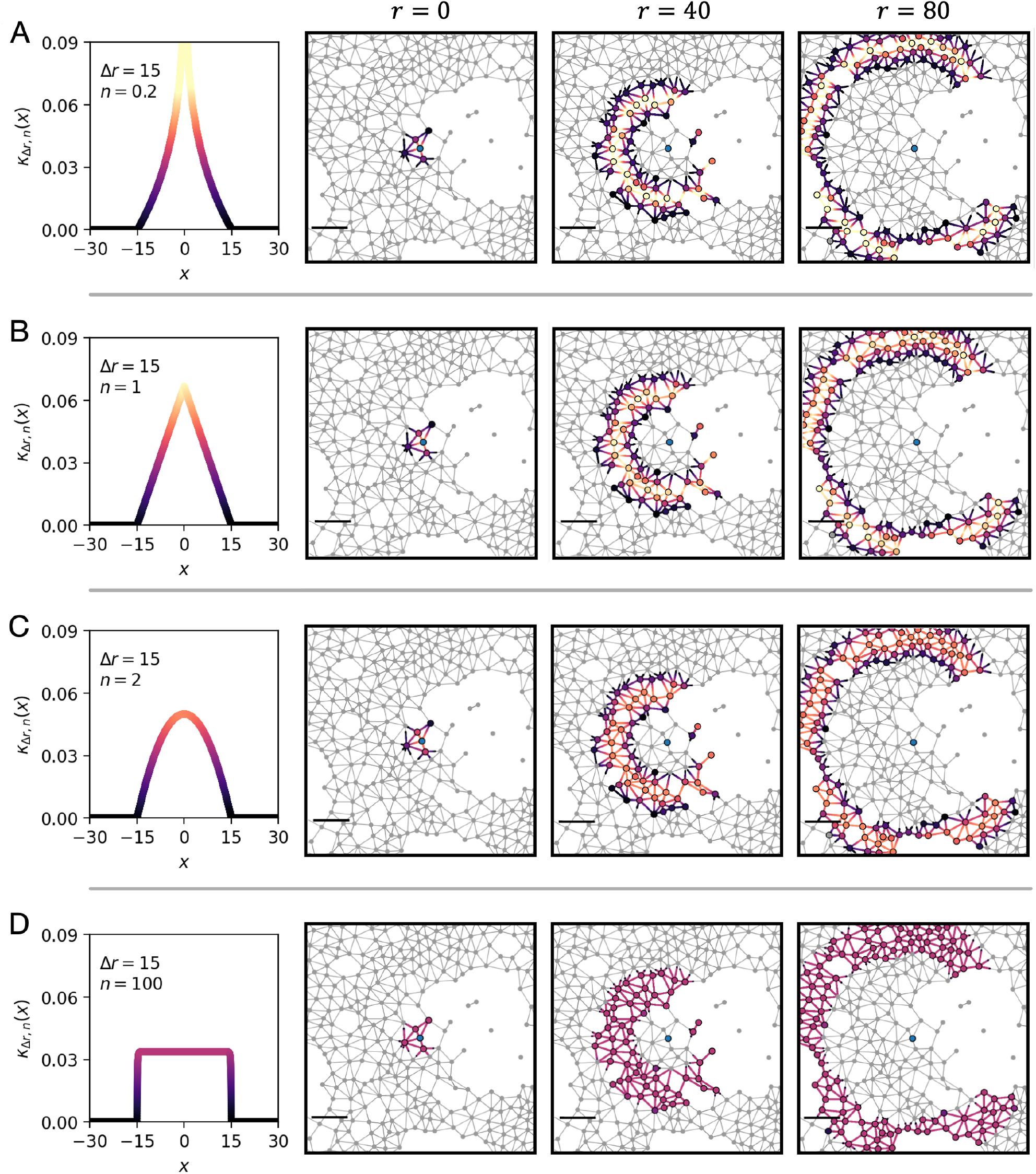
A range of polynomial kernels resolved on a spatial network. The polynomial kernel of finite support (Eq. 3) is shown with bandwidth Δ*r* = 15 and varying kernel shape parameter *n*, evaluated around an exemplar node *v*_*i*_ (central blue node) with other nodes coloured by kernel contribution at distances *r* = 0, 40, and 80. (A) *n* = 0.2, (B) *n* = 1, (B) *n* = 2 (Eponechnikov), and (C) *n* = 100. Scale bar is 50 distance units.

### 1.2 Evaluation of the polynomial kernel normalisation term, *L*_*i*_(*r*)

Given the polynomial kernel *κ*_Δ*r,n*_(*x*) (Eq. (3)), the accessible kernel length about a node *v*_*i*_, denoted *L*_*i*_(*r*) (Eq. (2)), can be computed explicitly by integrating the kernel over the network domain.

Let *d*_*ij*_ = *d*_sp_(*v*_*i*_, *v*_*j*_) denote the weighted shortest-path distance between nodes *v*_*i*_ and *v*_*j*_. The contribution to *L*_*i*_(*r*) from an edge *e*_*st*_ ∈ *E* of length *w*_*st*_ is obtained by parameterising points along the edge via *x* ∈ [0, *w*_*st*_], yielding

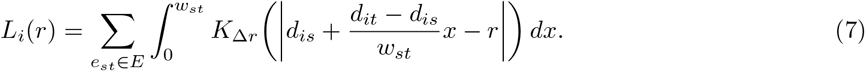

Substituting the explicit form of the kernel (Eq. (3)) gives

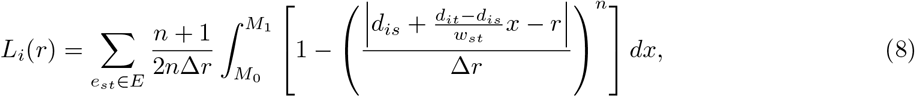

where the integration limits [*M*_0_, *M*_1_] correspond to the portion of the edge lying within the kernel support | · | *<* Δ*r*, given by

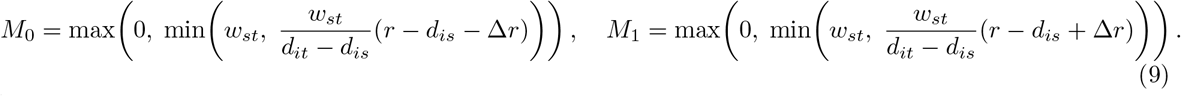

The integral separates into a constant term and a polynomial term,

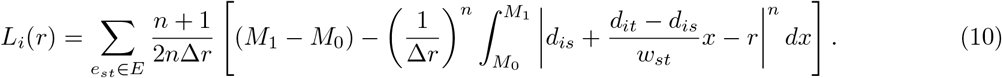

The integrand changes sign at

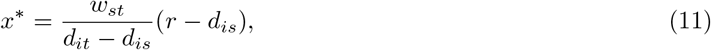

allowing the integral to be recast at with the intermediate point

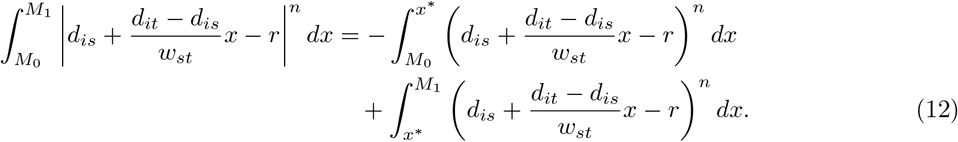

Solving Eq. (12) and substituting it into Eq. (10), we obtain the explicit expression for the accessible length of the kernel of over an arbitrary weighted network

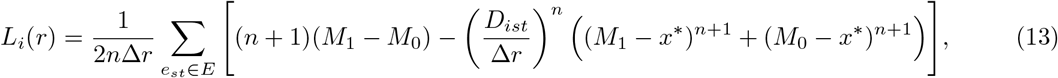

for 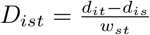, the weighted detour ratio. The explicit formulation of *L*_*i*_(*r*) in Eq. (13) provides an efficient means of evaluating the kernel normalisation term, as contributions from individual edges can be computed independently and vectorised across nodes in practical implementations.

### 1.3 Feature kernels for extended pair correlation functions

All pair correlation statistics considered in this work can be written in a common form by introducing feature kernels that act on node attributes. Let *G* = (*V, E*) be a spatial network with *N* = |*V* | nodes, let *d*_*ij*_ denote the weighted shortest-path distance between nodes *v*_*i*_ and *v*_*j*_, and let *L* = ∑_*i,j*_ *w*_*ij*_ denote the total network weighted length. As in the main text, let *L*_*i*_(*r*) denote the corresponding edge-correction term at distance *r* from node *v*_*i*_, and let *K*_Δ*r*_ be the spatial kernel defined in Eq. (3).

To allow for unmarked, categorical, and continuous data within a single framework, we define two generic feature functions,

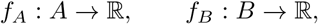

acting on node attributes 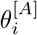 and 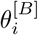. Their associated normalising constants are

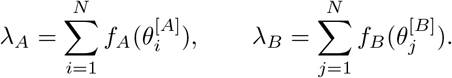

The corresponding general network pair correlation function may then be written as

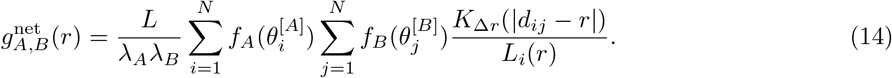

Specific choices of *f*_*A*_ and *f*_*B*_ recover the standard netPCF, the cross-netPCF, the weighted netPCF, and the cross-weighted netPCF [13].

#### Network pair correlation function (netPCF)

For an unmarked point pattern, all nodes are weighted equally, so that

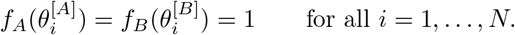

In this case, *λ*_*A*_ = *λ*_*B*_ = *N*, and Eq. (14) reduces to

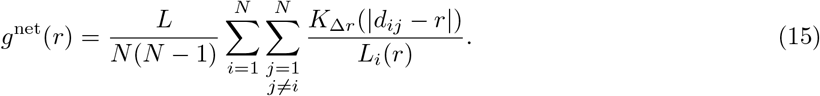

#### Network cross pair correlation function (cross-netPCF)

For two categorical marks, let 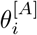 and 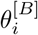 denote the categorical states at node *i*. For target categories *θ*_*A*_ ∈ *A* and *θ*_*B*_ ∈ *B*, define indicator feature functions

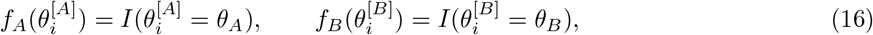

where *I*(·) is the indicator function. The corresponding normalising constants are

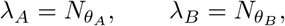

where 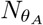 and 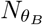 denote the number of nodes belonging to the respective target categories. The cross-netPCF is therefore

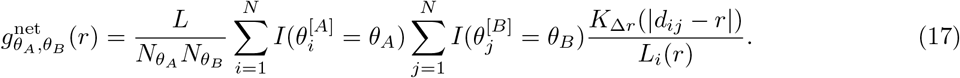

#### Weighted network pair correlation function (weighted-netPCF)

For a categorical mark and a continuous mark, let 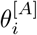 be categorical and 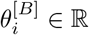 be continuous. For target category *θ*_*A*_ ∈ *A* and target continuous value *τ*_*B*_ ∈ ℝ with bandwidth Δ*τ*_*B*_, define

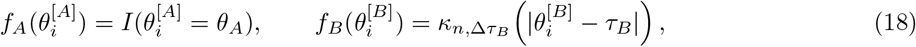

where 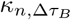 is the polynomial mark kernel of shape parameter *n*. The continuous-mark mass is

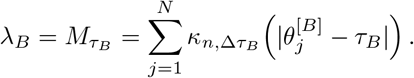

The weighted netPCF is then

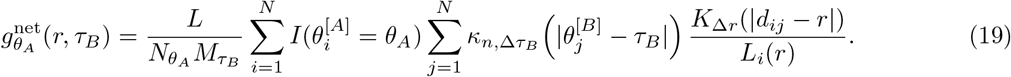

#### Cross-weighted network pair correlation function (cross-weighted-netPCF)

For two continuous marks, let 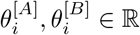 For target values *τ*_*A*_, *τ*_*B*_ ∈ ℝ, define

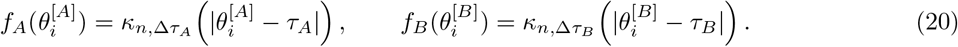

The corresponding normalising constants are

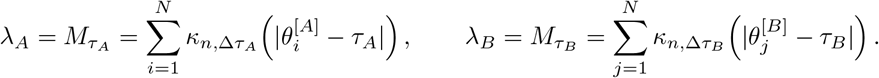

The cross-weighted netPCF is then

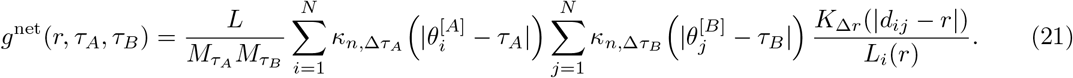

This statistic measures spatial association between two continuous marks at prescribed values *τ*_*A*_ and *τ*_*B*_. The single-mark continuous case is recovered by setting 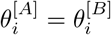.

### 1.4 Tiling a spatial network: Compact and volume-balanced partitions

In this section, we introduce a method for partitioning spatial networks into regions that are approximately equal in volume and geometrically compact. Such partitions are required for spatial resampling procedures, including the estimation of uncertainty in correlation functions.

Existing approaches to partition graphs are typically framed in the context of community detection, where the objective is to identify groups of nodes that are more densely connected internally than externally [22]. Such modularity-based approaches and their algorithmic implementations (e.g. Louvain and Leiden [6, 56]), are designed to recover intrinsic structure in the graph. However, they are not appropriate for the present setting, where the goal is to divide the spatial domain into contiguous regions of comparable size rather than to identify communities. In particular, modularity-based methods do not explicitly control for region size and can produce highly imbalanced partitions, which is undesirable for spatial resampling (Fig. 11).

**Fig 11.**
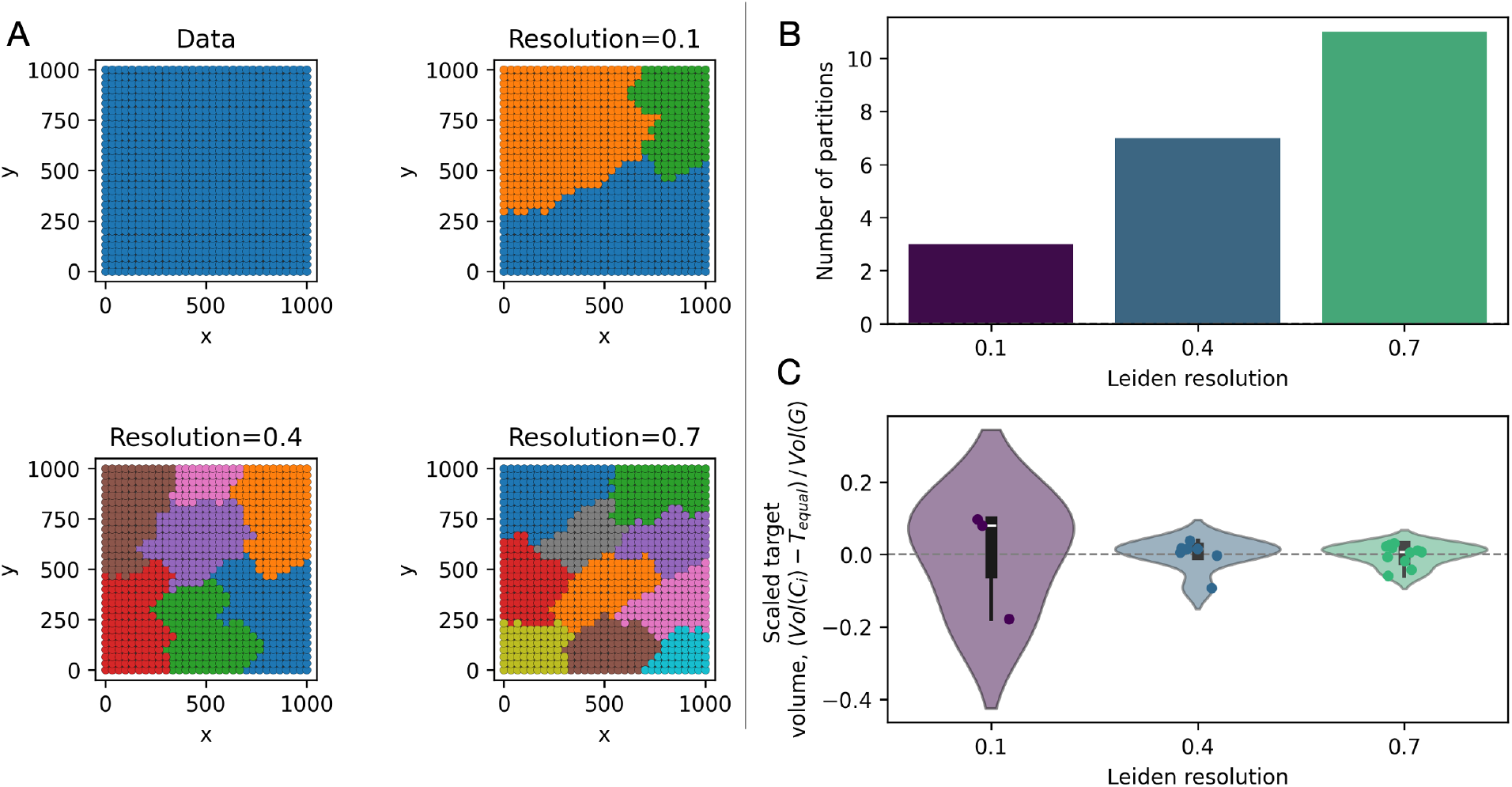
Modularity-based partitions are insufficient for spatial network tiling. (A) A spatial network defined by an orthogonal regular lattice of vertices in a 1000 × 1000 region. Plots are shown for all data (no partitions), and nodes colored by partitions generated from the Leiden algorithm for increasing resolution parameters. (B) Bar plot of the number of generated clusters produced from the Leiden algorithm at increasing resolutions. (C) Violin plots of the difference of partition volume, *V ol*(*C*_*i*_) and the volume produced from a regular tiling, *T*_*equal*_ = *V ol*(*C*_*i*_)*/k* for *k* the number of clusters. Results are shown for increasing resolutions of the Leiden algorithm [56].

Nevertheless, community detection algorithms provide useful algorithmic primitives, particularly in terms of efficient local refinement procedures and their ability to respect topological constraints such as connectivity. In this work, we adapt these ideas, specifically, Leiden-style node refinement [56], to construct node partitions that are connected, geometrically compact, and approximately volume-balanced.

We emphasise that we use the term *partition* rather than *community* to reflect this distinction. Spatial networks are typically sparse and represent an embedding of a geometric domain, rather than an abstract relational structure. As such, the objective here is to subdivide the domain into smaller, contiguous regions that respect the underlying geometry, rather than to detect densely connected subgraphs.

We below describe the algorithm for constructing these partitions and then validate the method a range of synthetic spatial networks with diverse geometries.

#### Algorithm outline

Let *G* = (*V, E*) be an undirected graph with non-negative edge weights *w*_*uv*_, interpreted as distances. The goal is to partition the node set *V* into *k* connected partitions {*C*_1_, …, *C*_*k*_ } such that each community is both geometrically compact (in the graph metric) and approximately equal in volume.

The *volume* of a partition *C*_*j*_ is defined as the total edge weight incident to its nodes,

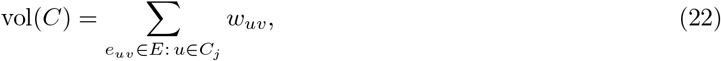

so that boundary edges contribute to the volume of both adjacent partitions.

Each partition *C*_*j*_ is represented by a *medoid* node *m*_*j*_ ∈ *C*_*j*_. Let *d*(*m*_*j*_, *u*) denote the shortest-path distance between *m*_*j*_ and *u* in *G*. The partition is obtained by minimising the objective function

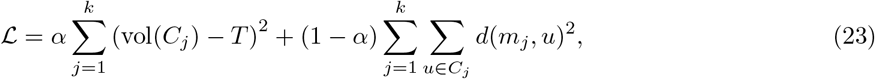

where *T* is a target partition volume and *α* ∈ [0, 1] controls the trade-off between volume balance and compactness.

#### Initialisation

If the number of partitions *k* is specified, *k* seed nodes are selected using a farthest-first traversal under the shortest-path metric. Nodes are then assigned to their nearest seed via a multi-source Dijkstra search, yielding a graph Voronoi partition. If the target volume *T* is not specified, it is initialised as the mean partitions volume under this partition, *T* = vol(*G*)*/k*. Any disconnected community is split into its connected components to ensure topological coherence.

#### Medoid estimation

For a fixed partition, medoids are approximated independently for each community. A subset of nodes is sampled, and the node minimizing the sum of shortest-path distances to all other nodes in the community is selected. This provides an efficient approximation to the graph *k*-median problem.

#### Local refinement

The partition is refined via greedy local node moves. All nodes are visited in random order and considered for reassignment only to adjacent communities. For each candidate move, the exact change in the objective function is computed, accounting for both the compactness term and the volume penalty. Moves that would disconnect the source partition are disallowed, and a node is reassigned only if the move yields a strict decrease in ℒ.

Medoids are recomputed after each round of node updates, and refinement passes are repeated until convergence. Convergence is declared under either of the following conditions: (i) the change in the objective function between successive iterations is below a threshold, |ℒ_*i*_ − ℒ_*i*+1_| *< δ*, for *δ* ≪ 1, or (ii) no nodes are reassigned during a refinement pass.

#### Computational complexity

The dominant cost arises from repeated shortest-path computations from community medoids. Let *n* = |*V*|, *m* = |*E*|, and *k* be the number of communities. In the worst case, each iteration scales as *O*(*k m* log *n*), although caching, medoid sampling, and the locality of node updates substantially reduce runtime in practice.

#### Summary

The proposed method combines graph Voronoi initialisation, medoid-based compactness optimisation, and Leiden-style greedy refinement, while enforcing a soft quadratic constraint on partition volume. Unlike modularity-based approaches, the objective is explicitly spatial metric-aware and allows direct control over partitions volume. Psuedo-code for the algorithm is provided below.

##### Algorithm 1 Pseudo-code: compact and volume-balanced partitions

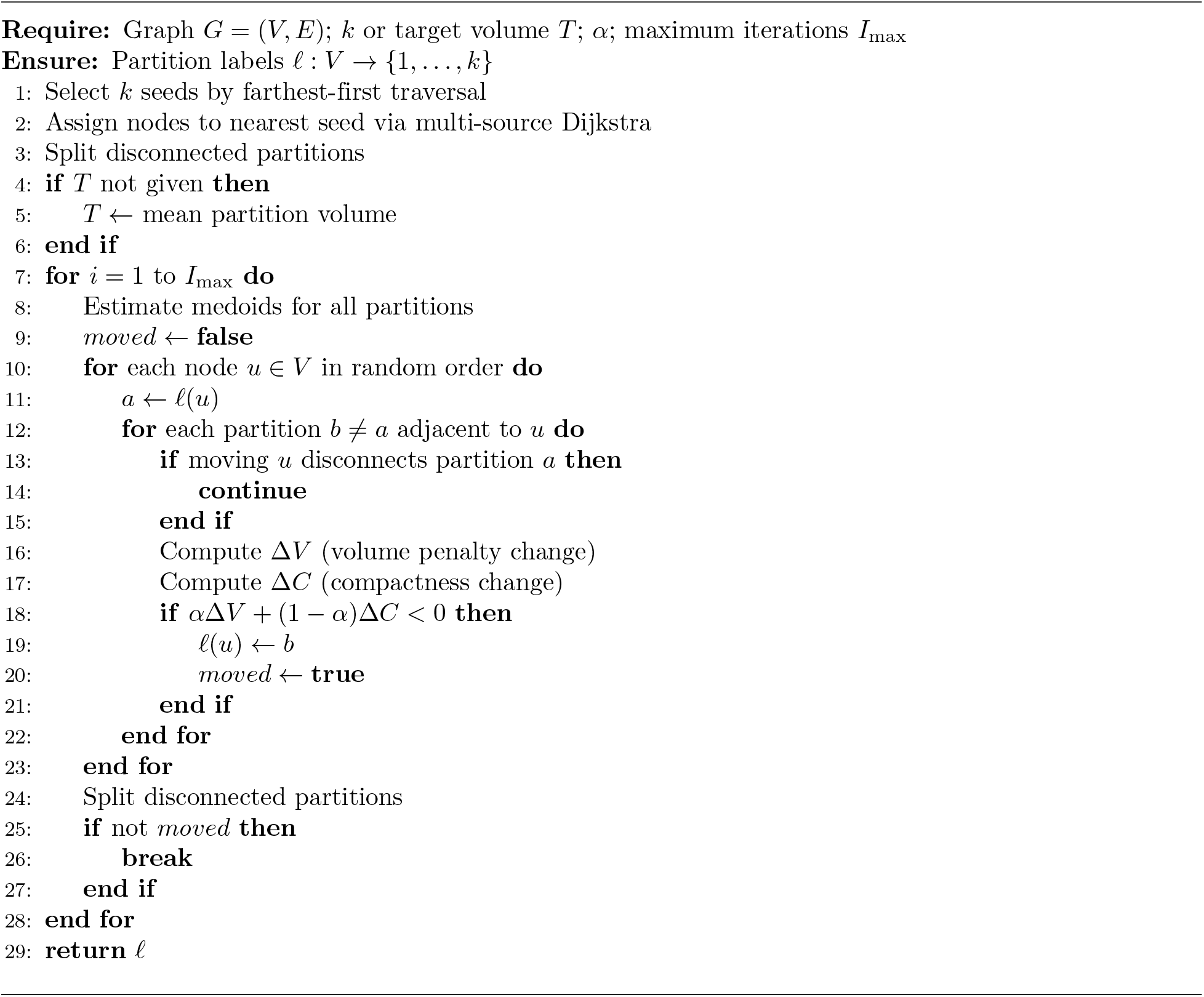

#### Validation

We validate the partitioning algorithm on a suite of synthetic spatial networks with increasing geometric complexity (Fig. 12). These include a regular orthogonal lattice (Grid), a homogeneous Poisson point process (Poisson), a Poisson process with excluded regions (Poisson with holes), a Poisson process with excluded regions and superimposed clustering (Poisson with holes and clusters), and a curved spiral domain.

**Fig 12.**
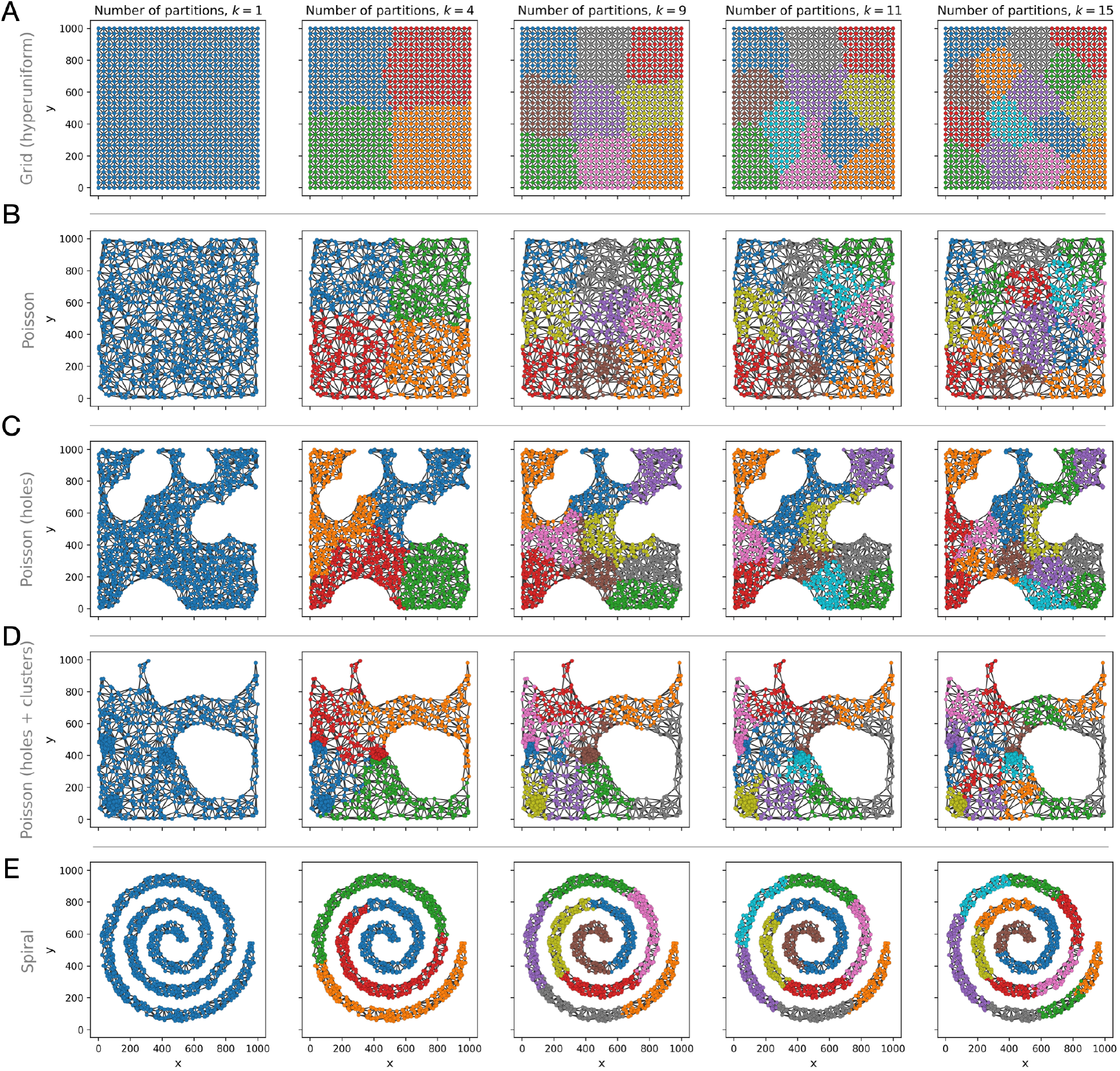
Compact and volume-preserving partitions of spatial networks. Representative partitions of synthetic spatial networks with nodes coloured by community assignment for increasing numbers of partitions (*k* = 4, 9, 11, 15). Networks are constructed from a range of spatial domains: (A) regular orthogonal lattice (Grid), (B) homogeneous Poisson point process (Poisson), (C) Poisson point process with excluded regions (Poisson with holes), (D) Poisson point process with excluded regions and superimposed clustering (Poisson with holes and clusters), and (E) curved domain defined by a Poisson point process along a spiral (Spiral).

For each network, partitions are computed for *k* = 4, 9, 11, 15 using *α* = 0.6. The resulting partitions are shown in Fig. 12, where nodes are coloured by partition assignment. Across all datasets and values of *k*, the algorithm produces the prescribed number of spatially contiguous and geometrically coherent partitions. For geometrically simple domains (Grid and Poisson), the resulting partitions closely resemble regular tilings, analogous to those obtained via sliding-window approaches (Fig. 12A–B). As domain complexity increases (Poisson with holes, clustered domains, and spiral geometries), partitions adapt to the underlying structure while remaining connected and similar in volume (Fig. 12C–E). In particular, regions of higher local edge density (e.g. clustered areas) are assigned fewer nodes, reflecting the use of edge-weight-based volume and indicating that the method accounts for spatial heterogeneity in the domain.

#### Convergence

The optimisation objective ℒ converges rapidly across all datasets and values of *k*, typically stabilising within five iterations (Fig. 13A).

**Fig 13.**
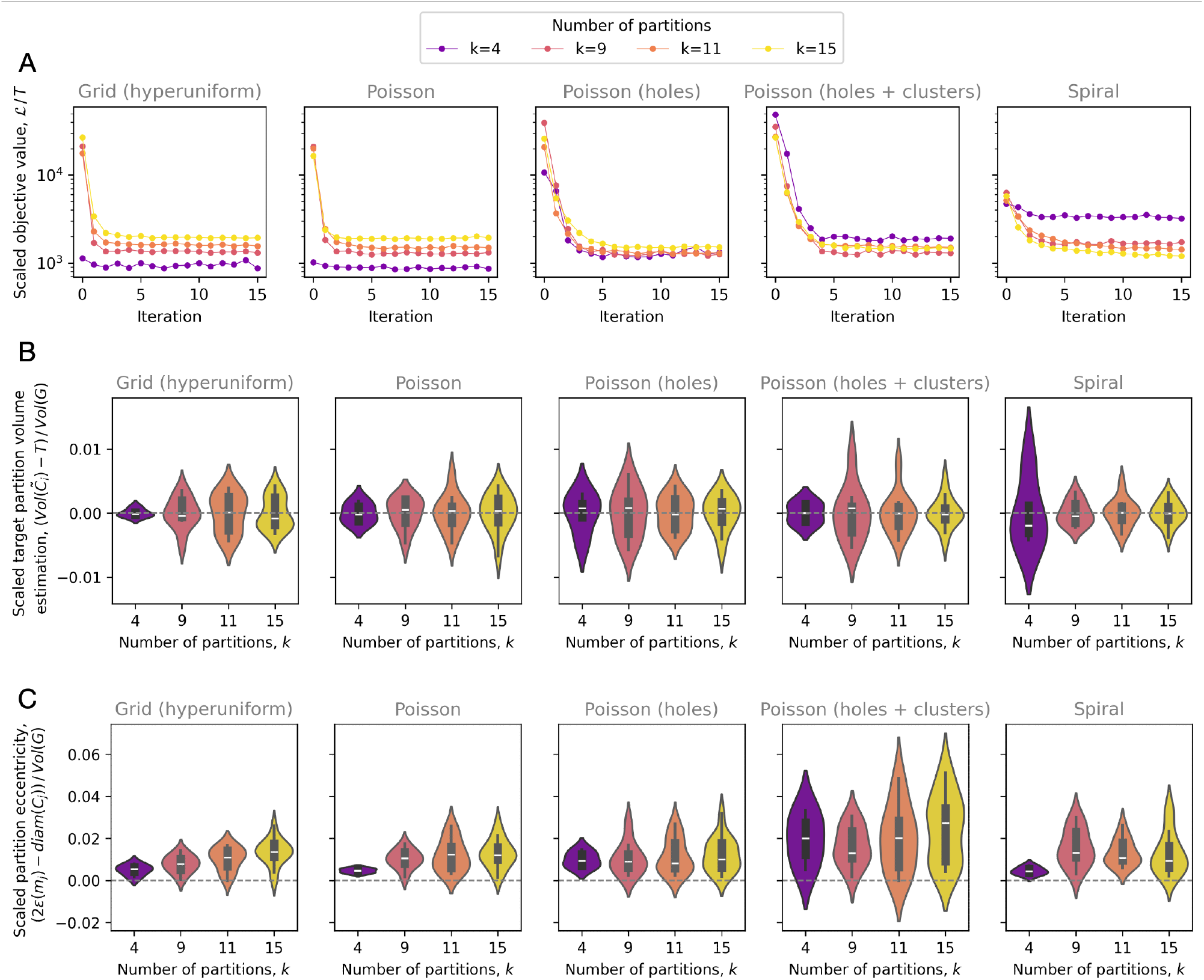
Quantitative evaluation of compact and volume-preserving partitions. (A) Convergence of the objective function ℒ (scaled by the target partition volume *T* ) across optimisation iterations, shown for all spatial networks and numbers of partitions (*k*) on log-scale. (B) Distribution of final partition volumes 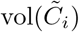 relative to the target volume *T*, normalised by the total network volume vol(*G*). Violin plots are shown for each value of *k* = 4, 9, 11, 15 across all spatial networks. (C) Distribution of partition compactness, quantified by the eccentricity *ϵ*(*m*_*j*_) relative to the diameter diam(*C*_*j*_), normalised by the total network volume vol(*G*). Violin plots are shown for each value of *k* = 4, 9, 11, 15 across all spatial networks.

#### Volume preservation

To assess volume balance, we compute the effective volume of each partition,

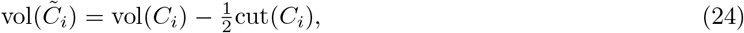

accounting for both internal edges and shared boundary contributions. As shown in Fig. 13B, partition volumes are tightly distributed around the target volume *T* with low variance across all datasets and values of *k*. While variance increases slightly for more complex domains (e.g. Poisson with holes and clusters), the deviation remains small, demonstrating effective volume preservation.

#### Compactness

To quantify geometric compactness, we compute the eccentricity of each partition, *ϵ*(*m*_*j*_), defined as the maximum shortest-path distance from the medoid *m*_*j*_ to nodes in *C*_*j*_, and compare this to the partition diameter diam(*C*_*j*_). For perfectly compact (e.g. convex or isotropic) regions, 2*ϵ*(*m*_*j*_) − diam(*C*_*j*_) = 0. Deviations from zero indicate asymmetry or elongation of the partition. Across all datasets, partitions exhibit low positive deviations, indicating good compactness with slight geometric asymmetries (Fig. 13C). As expected, variability is higher in geometrically heterogeneous domains, but overall compactness remains well-controlled.

#### Summary

These results demonstrate that the proposed algorithm produces spatially contiguous, volume-balanced, and geometrically compact partitions across a range of network geometries. This provides a robust and computationally efficient analogue of tiling for arbitrary weighted networks, (and in particular spatial networks), suitable for downstream spatial resampling and uncertainty estimation.

### 1.5 Definition of the Euclidean pair correlation function

In this section, we briefly define the pair correlation function (PCF) in Euclidean space, which we refer to as the *Euclidean PCF* throughout this study.

Let 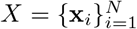 denote a set of *N* points observed within a spatial domain Ω ⊂ ℝ^2^ with total area *A*. Under complete spatial randomness (CSR), and using the limiting top-hat form of the polynomial kernel as *n* → ∞, the *Euclidean PCF* is defined as

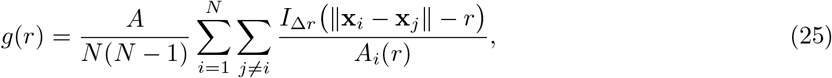

where ∥**x**_*i*_ − **x**_*j*_∥ denotes the Euclidean distance between points **x**_*i*_ and **x**_*j*_. The indicator function *I*_Δ*r*_(· ) selects pairs separated by distances within a band of width 2Δ*r* centred at *r*, given by

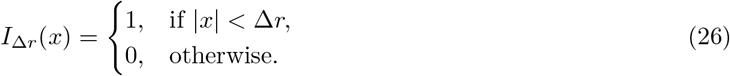

The normalisation term *A*_*i*_(*r*) accounts for boundary effects and corresponds to the area of the annulus of width 2Δ*r* centred at distance *r* from point **x**_*i*_, truncated by the domain Ω.

Euclidean PCFs are computed using the implementation provided in the `MuSpAn` package (v1.2.4) [15], via the `muspan.spatial_statistics.cross_pair_correlation_function` method.

### 1.6 Bandwidth lower bounds for sufficient kernel coverage on data-driven spatial networks to mitigate domain discretisation effects

When a spatial network is constructed directly from observed point data, its geometry may influence subsequent estimates of spatial association. This is particularly important for network pair correlation functions, where the same set of points may define the spatial domain and provide the locations on which labels or marks are analysed. In this setting, insufficient kernel bandwidth may introduce an artificial local exclusion effect: if the kernel support is too narrow relative to the spacing between adjacent network nodes, then nearby distances are undersampled by construction, producing apparent short-range negative correlation even under complete spatial randomness.

We introduce a bandwidth-selection criterion that ensures discretisation of the spatial embedding does not introduce misleading correlations into the estimated PCF. We assume that the point pattern used to define the network has approximately homogeneous density, so that large-scale variation in node density is not the primary source of bias. The relevant geometric constraint is instead local coverage: the spatial kernel should place sufficient mass over the characteristic distance separating neighbouring nodes.

Let ⟨*l*⟩ denote a characteristic adjacent-node length scale. In practice, this can be defined from the network itself, for example as the mean edge length, or more conservatively as an upper quantile of the edge-length distribution, such as the 95% quantile. Alternatively, when relevant prior information is available, ⟨*l*⟩ may be chosen to reflect an application-specific physical length scale, such as a typical cell diameter. A necessary condition for avoiding local exclusion

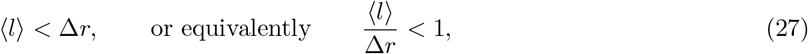

where Δ*r* is the kernel bandwidth. However, this condition alone is not sufficient, as the amount of kernel mass contained below ⟨*l*⟩ depends on the kernel shape (Fig. 14). For polynomial kernels with small exponent *n*, relatively little mass may lie near the boundary of the kernel support, so exclusion effects can persist even when ⟨*l*⟩ is slightly smaller than Δ*r*.

**Fig 14.**
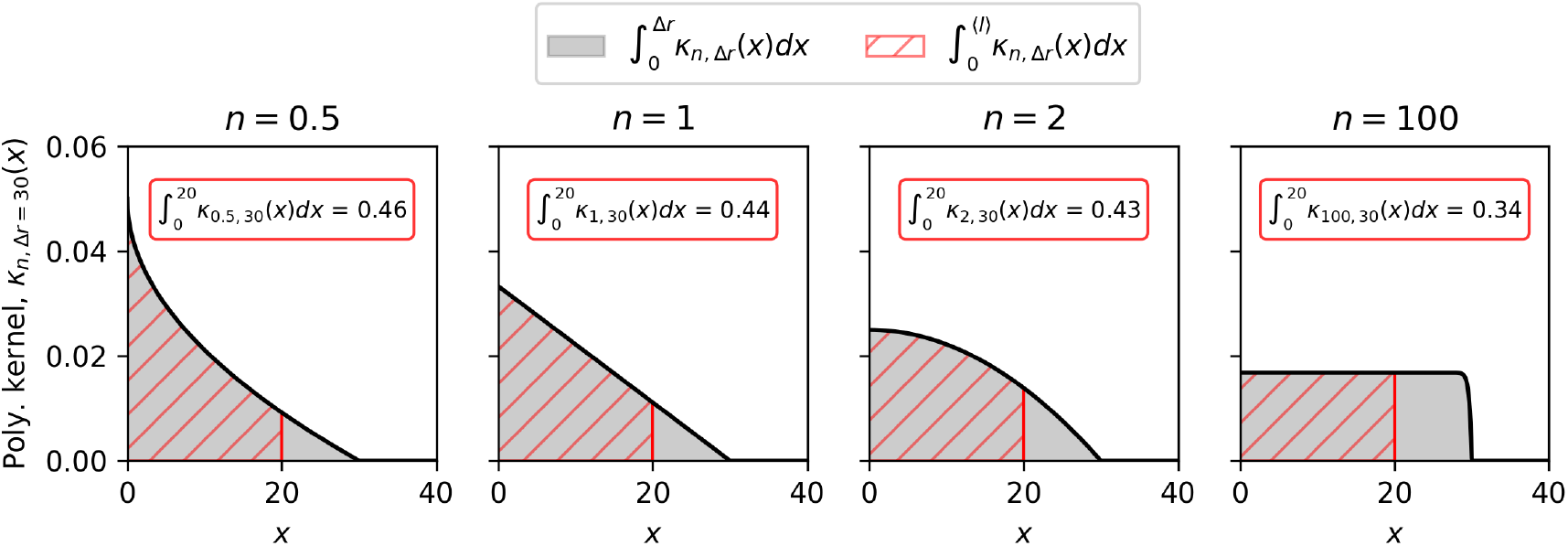
Polynomial kernel coverage at a characteristic length scales. Polynomial kernels (Eq. (3)) are shown for a fixed bandwidth Δ*r* = 30 and kernel exponents *n* = 0.5, 1, 2, and 100. Hatched regions indicate the proportion of kernel mass contained within a characteristic network length scale ⟨*l*⟩ = 20, with the corresponding integrated kernel mass reported above each curve.

We define sufficient coverage in terms of the proportion of kernel mass contained within the characteristic length scale of the spatial network. Let *α* ∈ (0, 1) be the desired fraction of one-sided kernel mass covered by distance ⟨*l*⟩. For the polynomial kernel, *κ*_*n*,Δ*r*_ (Eq. (3)), we impose

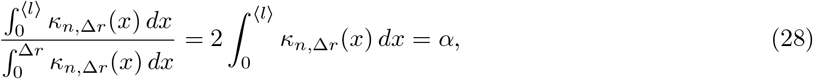

where the denominator follows from the normalisation of the symmetric polynomial kernel, as detailed in Section 1.1. Using Eq. (28), we seek to establish a relationship between ⟨*l*⟩ and Δ*r* for a desired *α* value. Evaluating the integrated kernel mass up to ⟨*l*⟩ gives

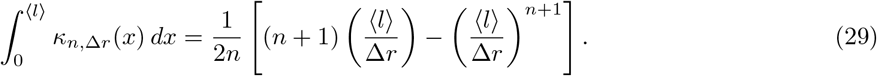

Substituting Eq. (29) into Eq. (28), and defining

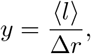

yields the polynomial equation

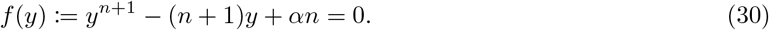

The admissible solution satisfies *y* ∈ (0, 1), corresponding to the necessary condition 0 *<* ⟨*l*⟩ *<* Δ*r*.

For all *n >* 0 and *α* ∈ (0, 1), Eq. (30) has a unique solution in (0, 1). Indeed,

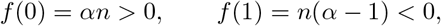

and

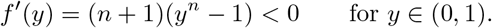

Thus *f* is strictly decreasing on (0, 1), and the Intermediate Value Theorem guarantees a unique root [55], denoted 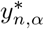. This root defines the minimum bandwidth required to achieve kernel mass coverage *α*:

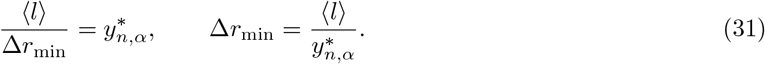

For selected values of *n*, closed-form solutions are available. When *n* = 1, Eq. 30 reduces to

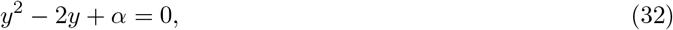

and the admissible root is

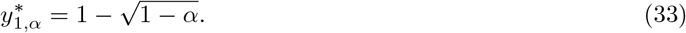

When *n* = 2, Eq. (30) becomes

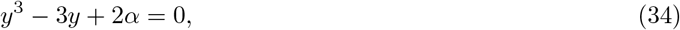

and the admissible root is

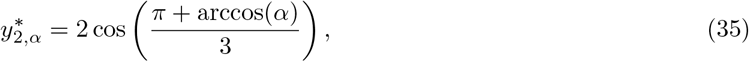

which lies in (0, 1) for all *α* ∈ (0, 1).

In addition, in the limit *n* → ∞, the polynomial kernel approaches the uniform kernel on [−Δ*r*, Δ*r*], with

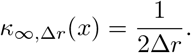

Namely, the polynomial solution Eq. (28) yields,

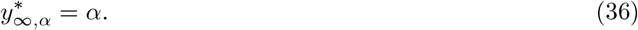

Although closed-form solutions of the polynomial (Eq. (30)) are not available for arbitrary *n >* 0, the reciprocal root can be accurately approximated by the ansatz

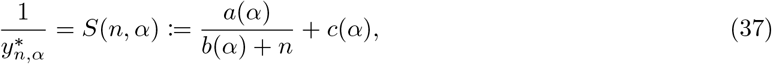

where the three parameterising functions, *a*(*α*), *b*(*α*), and *c*(*α*), are identified by matching the exact solutions at *n* = 1, *n* = 2, and *n* → ∞. This gives

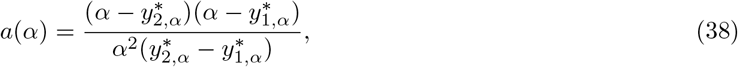

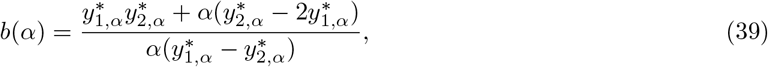

and

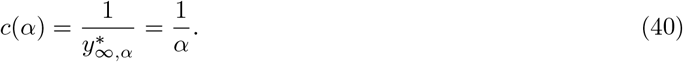

The approximation in Eq. (37) agrees closely with numerical solutions of Eq. (30) in the admissible parameter range (Fig. 15). Therefore, it provides an explicit approximation to the bandwidth scaling factor 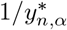 for any *α* ∈ (0, 1) and *n* ∈ ℝ_*>*0_.

**Fig 15.**
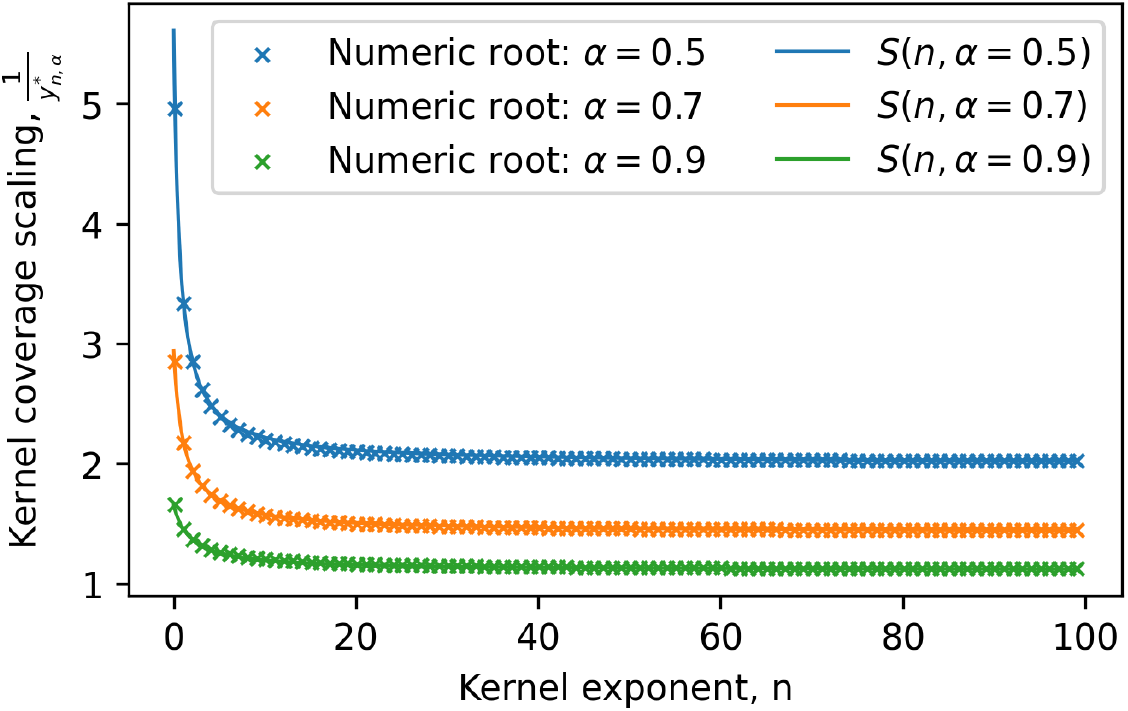
Approximation of polynomial kernel coverage solutions. Reciprocal numerical solutions of Eq. 30 are shown as scatter points as a function of the kernel exponent *n* for kernel coverage proportions *α* = 0.5, 0.7, and 0.9. Solid lines show the corresponding approximate solutions, *S*(*n, α*), for each value of *α*.

Consequently, the bandwidth lower bound can be written as

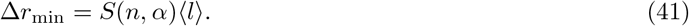

This expression gives a data-dependent lower bound on the kernel bandwidth, controlled by the kernel exponent *n*, the target coverage proportion *α*, and the characteristic network spacing ⟨*l*⟩ . Table 1 reports representative values of *S*(*n, α*), showing the multiplicative factor by which the characteristic edge length should be scaled to obtain the minimum bandwidth.

**Table 1.** Representative bandwidth scaling factors. Values of 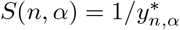, where Δ*r*_min_ = *S*(*n, α*)⟨*l*⟩. Lower values of *α* impose stronger kernel coverage requirements.

| Kernel exponent $n$ | $\alpha = 0.50$ | $\alpha = 0.75$ | $\alpha = 0.90$ | $\alpha = 1$ |
| --- | --- | --- | --- | --- |
| 0.5 | 4.03 | 2.19 | 1.54 | 1.00 |
| 1 | 3.41 | 2.00 | 1.46 | 1.00 |
| 2 | 2.88 | 1.79 | 1.37 | 1.00 |
| 5 | 2.41 | 1.57 | 1.25 | 1.00 |
| 10 | 2.21 | 1.46 | 1.20 | 1.00 |
| 100 | 2.02 | 1.35 | 1.12 | 1.00 |
| ∞ | 2.00 | 1.33 | 1.11 | 1.00 |

#### Sufficient kernel coverage for Poisson-derived spatial networks

For a homogeneous Poisson point process with intensity *ρ*, and in the absence of boundary effects, the characteristic spacing between neighbouring nodes scales as ⟨*l*⟩ ∝ *ρ*^−1*/*2^ [7]. Therefore,

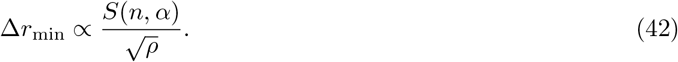

Thus, as *ρ* → ∞, the network discretisation converges locally to the Euclidean domain and the lower bound tends to zero, namely, spatial domain discretisation will not influence observed spatial correlations. In dense spatial embeddings, the kernel bandwidth is therefore no longer constrained by node separation. In sparse embeddings, however, Eq. (41) provides a sufficient safeguard against potential geometry-induced local exclusion.

To illustrate the sufficient kernel coverage criterion, we simulated a homogeneous Poisson point process with intensity *ρ* = 0.003, constructed the corresponding spatial network (Fig. 16A), and estimated the network PCF using an Epanechnikov kernel (*n* = 2). We defined the characteristic adjacent-node length as the mean edge length, giving ⟨*l*⟩ = 21.19. For a target coverage proportion of *α* = 0.75, Table 1 gives a bandwidth scaling factor of approximately *S*(2, 0.75) = 1.79. Therefore,

**Fig 16.**
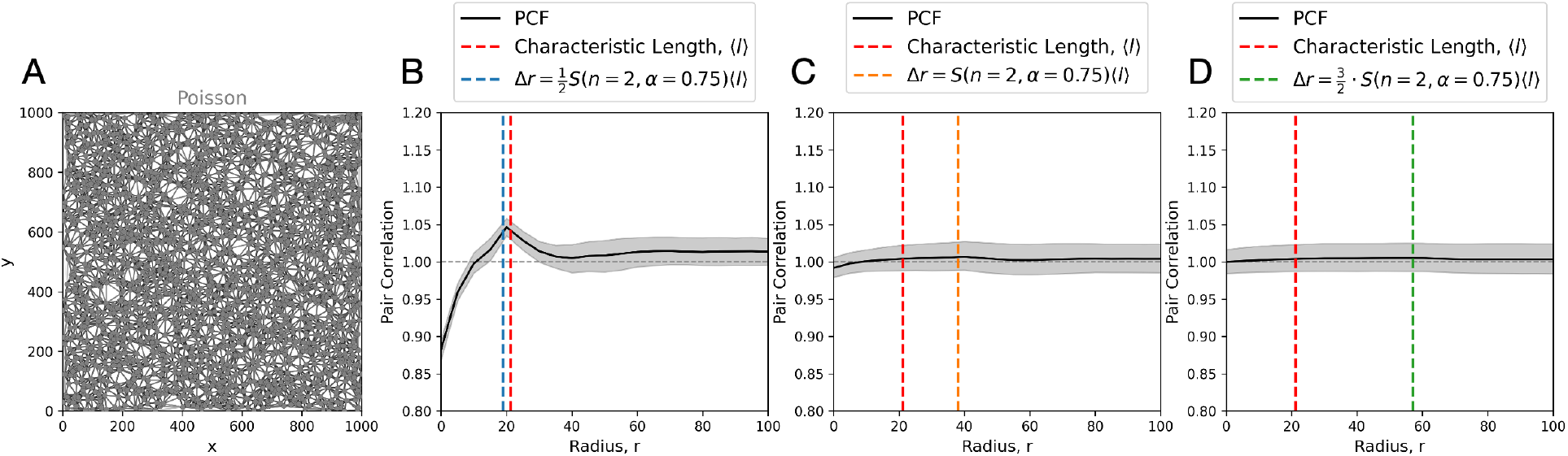
Mitigating discretisation effects through sufficient kernel coverage. A: Spatial network constructed from a point pattern realised from a homogeneous Poisson point process with intensity *ρ* = 0.003 in a 1000 × 1000 spatial domain. B–D: Estimated netPCF curves computed using increasing kernel bandwidths, Δ*r*, chosen relative to the lower bound in Eq. (41). The red dashed vertical line denotes the characteristic network length scale, ⟨*l*⟩, defined here as the mean edge length of the network in (A). The remaining dashed vertical line in each panel indicates the bandwidth used for that PCF computation, corresponding to 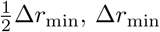, and 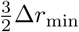 in B–D, respectively. Shaded regions indicate 95% confidence intervals around the estimated correlation.

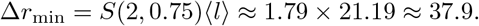

Thus, the sufficient-coverage lower bound corresponds to only 3.79% of the width of the spatial domain.

When Δ*r <* Δ*r*_min_, the estimated netPCF exhibited a short-range exclusion, followed by a transient co-localisation consistent with network edge length (Fig. 16B). These observations are removed when the bandwidth was increased to satisfy Δ*r* ≥ Δ*r*_min_ (Fig. 16C-D), recovering the expected complete spatial randomness baseline.

#### Summary

Local exclusion may be a genuine feature of the pair correlation in continuous domains, but it may be undesirable when the spatial network defined by data is intended only as a spatial support for estimating label or mark correlations. The sufficient kernel coverage criterion (Eq. (41)) provides a data-dependent lower bound on the kernel bandwidth, ensuring that geometric artefacts introduced by the sampled domain are not misinterpreted as biological or physical correlations of interest.

In the analyses of Results 1–4, background nodes were simulated with intensity *ρ* = 0.001. For the uniform-kernel limit, *n* → ∞, we selected Δ*r* = 35 (Results 1–2), and the Epanechnikov kernel, *n* = 2, we selected Δ*r* = 45 (Results 2–4), which both satisfy the sufficient kernel coverage lower bound (Eq. (41)) for the chosen coverage level *α* = 0.75.

### 1.7 Identifying patterns of continuous data over spatial networks

In Section, we showed that the cross-netPCF can identify and quantify spatial association between nodes carrying categorical marks. Here we consider the continuous analogue of that setting, in which nodes carry both categorical and continuous attributes. To analyse these data, we use the weighted-netPCF (Section 1.3), which measures how the spatial correlation structure changes when one population is filtered by a continuous mark kernel and compared against a reference population.

As in the categorical example, the weighted-netPCF detects the imposed structure across domains of increasing geometric complexity (Fig. 17). In the two-dimensional spiral, cylindrical surface, and three-dimensional volumetric examples, the continuous mark values vary systematically along the underlying geometry, while the categorical nodes are localised to specific regions of the domain. The resulting weighted-netPCF curves recover the imposed continuous pattern and show how the strength of association changes with the target mark value *τ*_*B*_. In this way, the method extends the cross-netPCF framework from discrete labels to continuous node attributes.

**Fig 17.**
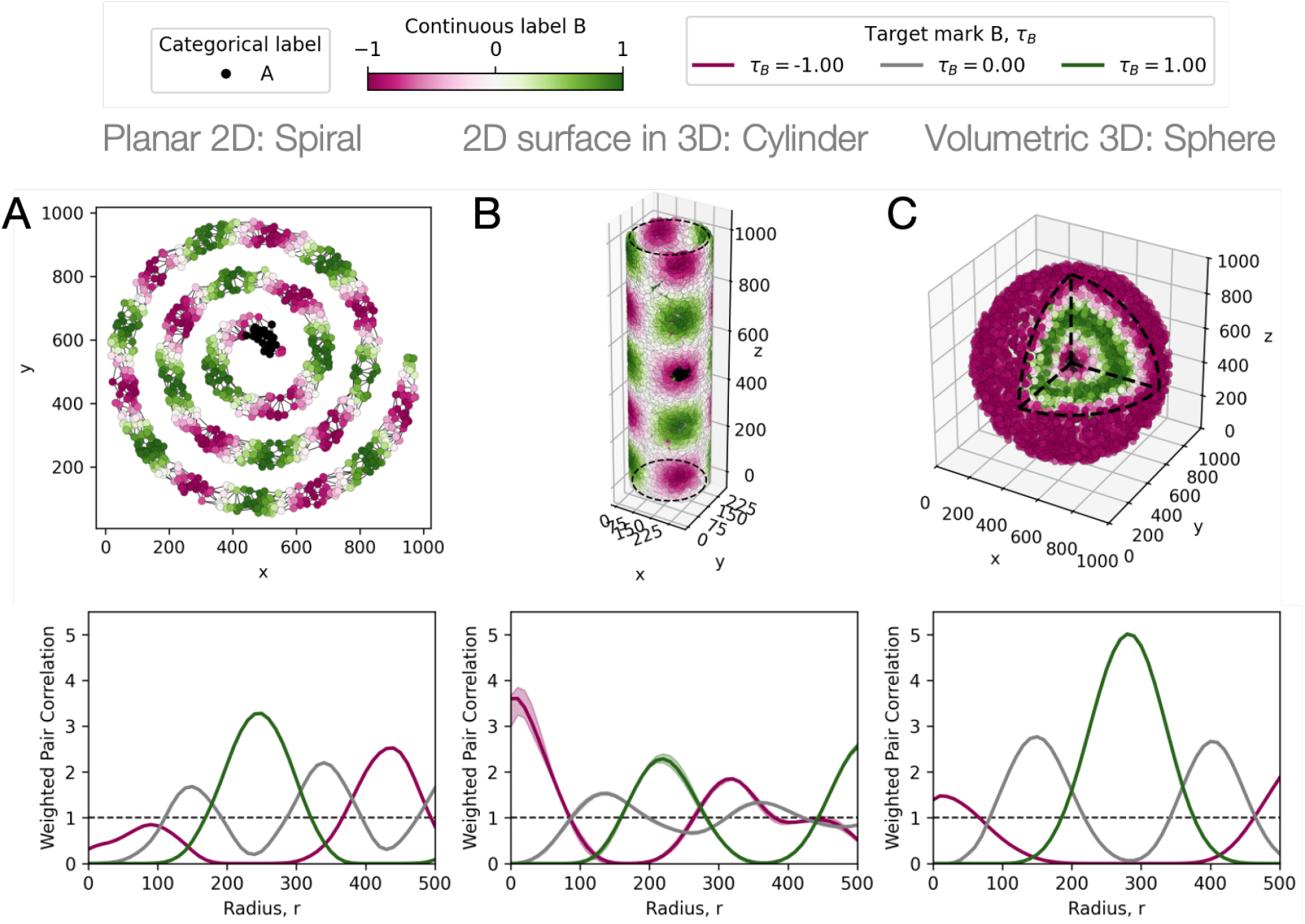
Quantifying continuous data on spatial networks. Synthetic point patterns with nodes assigned a combination of categorical (type A) and continuous (type B) attributes across domains of increasing geometric complexity. (A) Two-dimensional spiral domain, where points are generated from a Poisson point process along a curved trajectory. Nodes of type A (black) are located near the centre of the domain, while continuous values (type B) are assigned periodically as a function of distance along the spiral. (B) Two-dimensional surface embedded in three dimensions, defined by a hollow cylindrical domain (radius 137, height 1000), with points sampled from a Poisson point process over the surface. Continuous values (type B) are assigned periodically as a function of position on the cylinder, while nodes of type A are localised to a specific angular and axial region. (C) Three-dimensional volumetric domain, where points are generated from a Poisson point process within a sphere (radius 500). Nodes of type A are assigned to a central region (*<* 50 from the centre), while continuous values (type B) are assigned periodically as a function of radial distance. Weighted network-based PCF (netPCF) estimates between nodes of type A and nodes with continuous values (type B) are shown beneath each domain for target values *τ*_*B*_ = −1, 0, 1.

More generally, the cross-weighted-netPCF provides a corresponding measure of association between two continuous marks on the same network, allowing continuous–continuous spatial relationships to be quantified in the same framework (Section 1.3).

### 1.8 Extended information for robustness of netPCF to domain estimation

In main text, we present average Normalised Cross-Correlation (NCC) between baseline and the *in silico* perturbation experiments on Delaunay network structure for netPCF estimates. We present NCC values for all repeats in Fig. 18.

**Fig 18.**
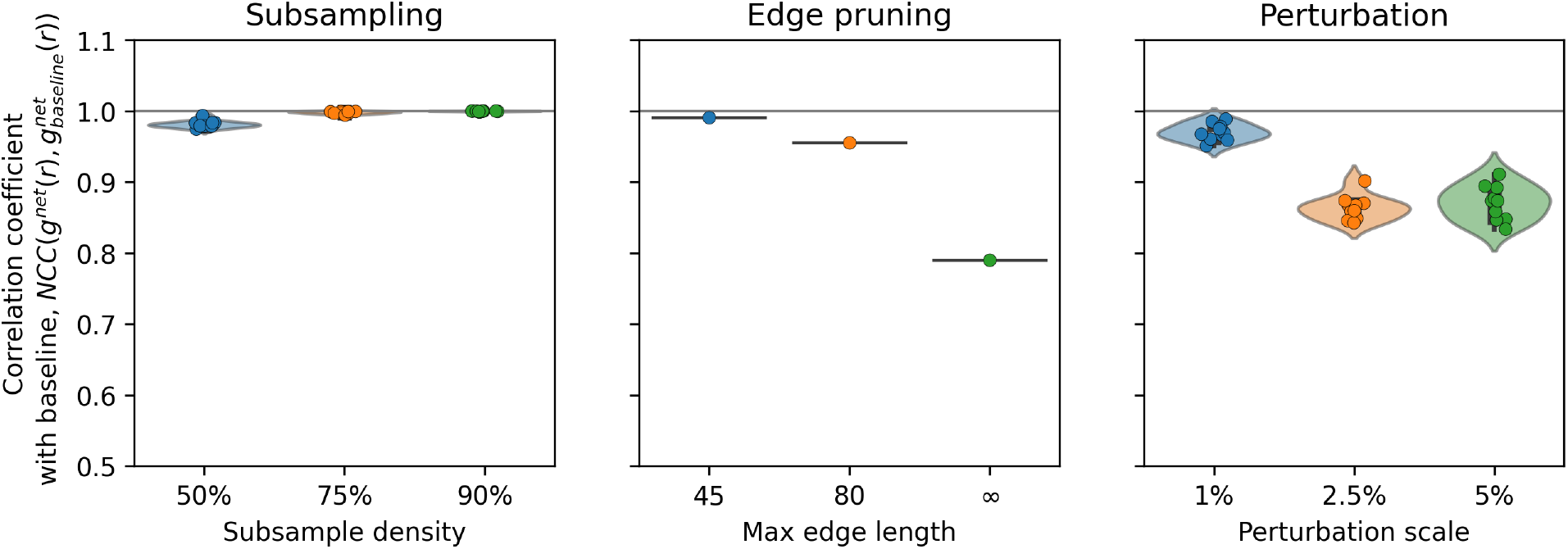
Quantitative comparison of network perturbations on netPCF estimates. Violin and jitter plots show normalised cross-correlation (NCC) values comparing perturbed netPCF estimates with the corresponding baseline estimate across the subsampling, edge-filtering, and spatial perturbation experiments.

### 1.9 Extended information for the 3D IMC Breast Carcinoma dataset and analysis

#### Parameters for computing netXPCF

The spatial networks representing the tissue domains were constructed as described in the Methods. For the breast carcinoma IMC case study, we selected an upper edge-length filtration threshold of *ϵ* = 40 µm to preserve the domain structure defined by morphologically extended cellular populations, including endothelial and fibroblast cells. This choice ensured sufficient local connectivity while maintaining the large-scale geometric organisation of the tissue.

For all netPCF and cross-netPCF estimations, we used a maximum interaction radius of 300 µm, chosen to provide coverage across the majority of the reconstructed tissue volume, whose total *z*-depth was 304 µm. Correlation functions were evaluated in increments of 5 µm.

Kernel parameters were selected using the lower-bound bandwidth estimates derived in Supplementary Information 1.6. Specifically, we employed a kernel shape parameter of *n* = 2, corresponding to an Epanechnikov kernel, with bandwidth Δ*r* = 45 µm for all 2D and 3D estimations. These parameters provided sufficient kernel support over the reconstructed spatial network while maintaining stable correlation estimates.

As a validation of the underlying spatial network representation, the unmarked netPCF was computed over all network nodes. This yielded *g*^net^(*r*) ≈ 1 for all *r <* 300 µm, indicating approximate spatial homogeneity of the reconstructed point cloud under the network metric (Fig. 19). Consequently, the marked cross-netPCF signals observed in the biological analyses are attributable to the spatial organisation of cellular populations rather than to large-scale inhomogeneity in the underlying spatial point density or inaccurate domain estimation via the chosen network.

**Fig 19.**
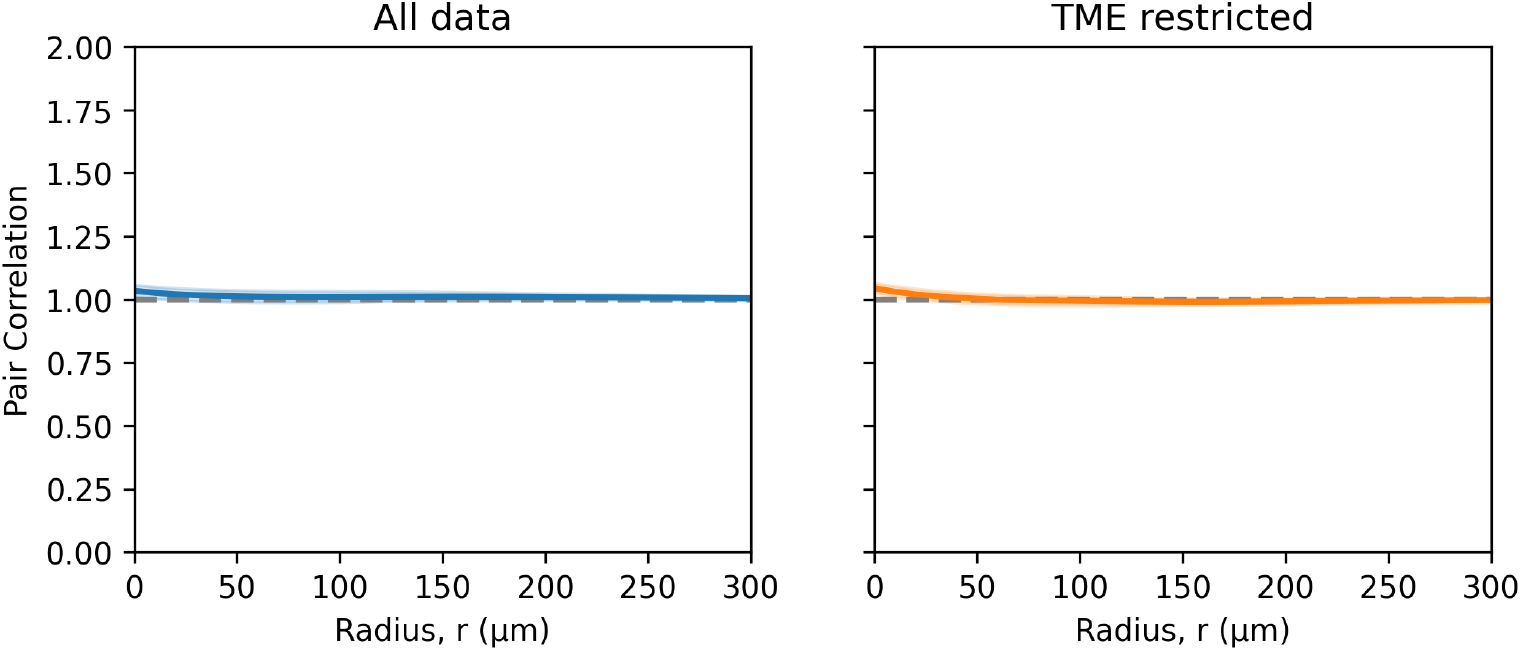
Pair correlation estimations of domain geometry: HER2+ breast carcinoma. netPCF estimates between all and TME only nodes of the HER2+ breast carcinoma data. Shaded regions denote 95% confidence intervals.

#### B and T cell spatial association

In addition to the endothelial–immune interactions presented in the main results, we computed cross-netPCF curves between B cells and T-cell subsets to assess immune–immune spatial organisation in the breast carcinoma IMC sample. Following the same analysis procedure, we compared full-domain and tumour microenvironment (TME)-restricted estimates in both 2D serial sections and the fully resolved 3D sample.

In 2D, the sparsity of B cells and T-cell populations results in highly variable cross-netPCF estimates. Although the curves show some evidence of longer-range exclusion between B cells and T cells, short-range associations are unstable and differences between full-domain and TME-restricted estimates are inconclusive (Fig. 20A). Thus, for these immune populations, individual 2D sections do not provide sufficient sampling density to robustly resolve local spatial interactions.

**Fig 20.**
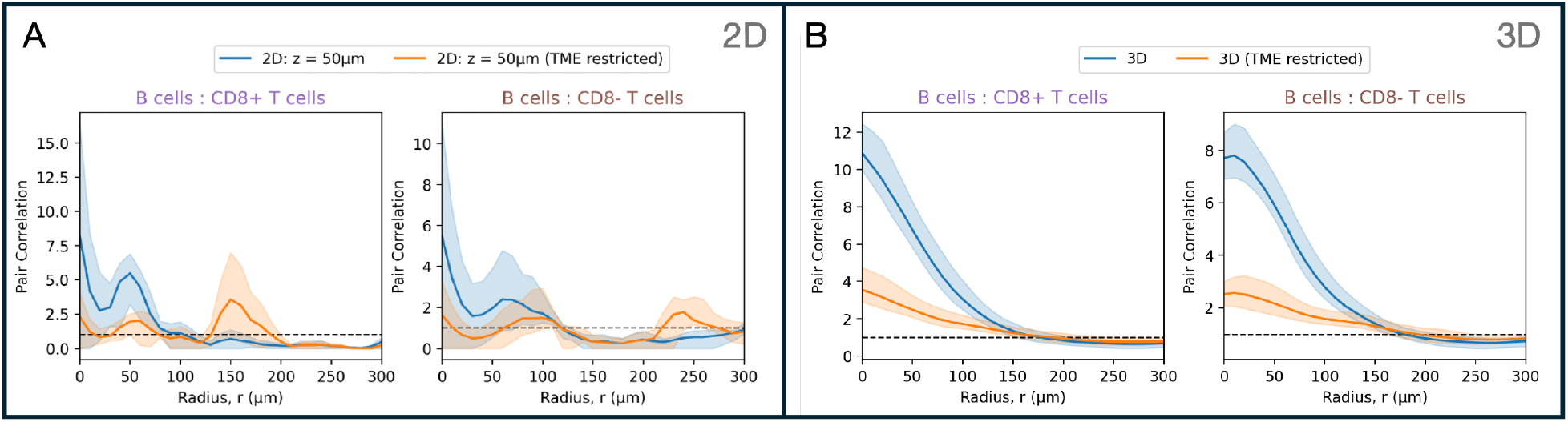
Immune–immune spatial interactions in a fully resolved breast carcinoma tumour microenvironment. Cross-netPCF curves quantify pairwise spatial associations between B cells and CD8+ or CD8-T cells in the breast carcinoma IMC sample. Estimates are shown for both the full tissue domain and the tumour microenvironment-restricted domain. (A) Representative 2D serial section at 50 µm depth from the IMC breast carcinoma sample. (B) Fully resolved 3D reconstruction of the IMC breast carcinoma sample. Shaded regions denote 95% confidence intervals.

In contrast, the 3D analysis shows clear qualitative agreement between the full-domain and TME-restricted estimates, with elevated short-range correlations between B cells and both CD8+ and CD8-T cells (Fig. 20B). This indicates robust short-range co-localisation and long-range exclusion of B and T cells that persists after restricting the analysis to the TME. Unlike the endothelial–immune interactions described in the main text, which were reduced under TME restriction, the persistence of B–T cell correlations suggests a domain-independent aggregation of these immune populations within stromal regions of the breast carcinoma sample.

These results further demonstrate the value of fully resolved 3D spatial data. The 2D serial section suggests a noisy and ambiguous interaction structure, whereas the 3D reconstruction resolves a consistent pattern of local B–T cell co-localisation. This contrast highlights how analyses based on individual 2D sections may produce incomplete or misleading interpretations when applied to sparse cell populations in heterogeneous tissue geometries.

#### Additional Endothelial–immune analysis in 2D serial sections

In the main text, we presented endothelial–immune cross-netPCF analysis for a representative 2D serial section at a depth of 50 µm. Here, we provide additional serial sections from the breast carcinoma IMC sample, each corresponding to a 4 µm section thickness, to illustrate the substantial spatial heterogeneity present throughout the tissue volume (Fig. 21). Although these sections remain a subset of the full dataset, they demonstrate the variability in endothelial and immune cell organisation across different depths of the sample. Notably, not all 2D planes contained sufficient representation of both endothelial and immune populations to support informative cross-netPCF estimation, further highlighting the limitations of analysing sparse cellular populations using individual 2D sections alone.

**Fig 21.**
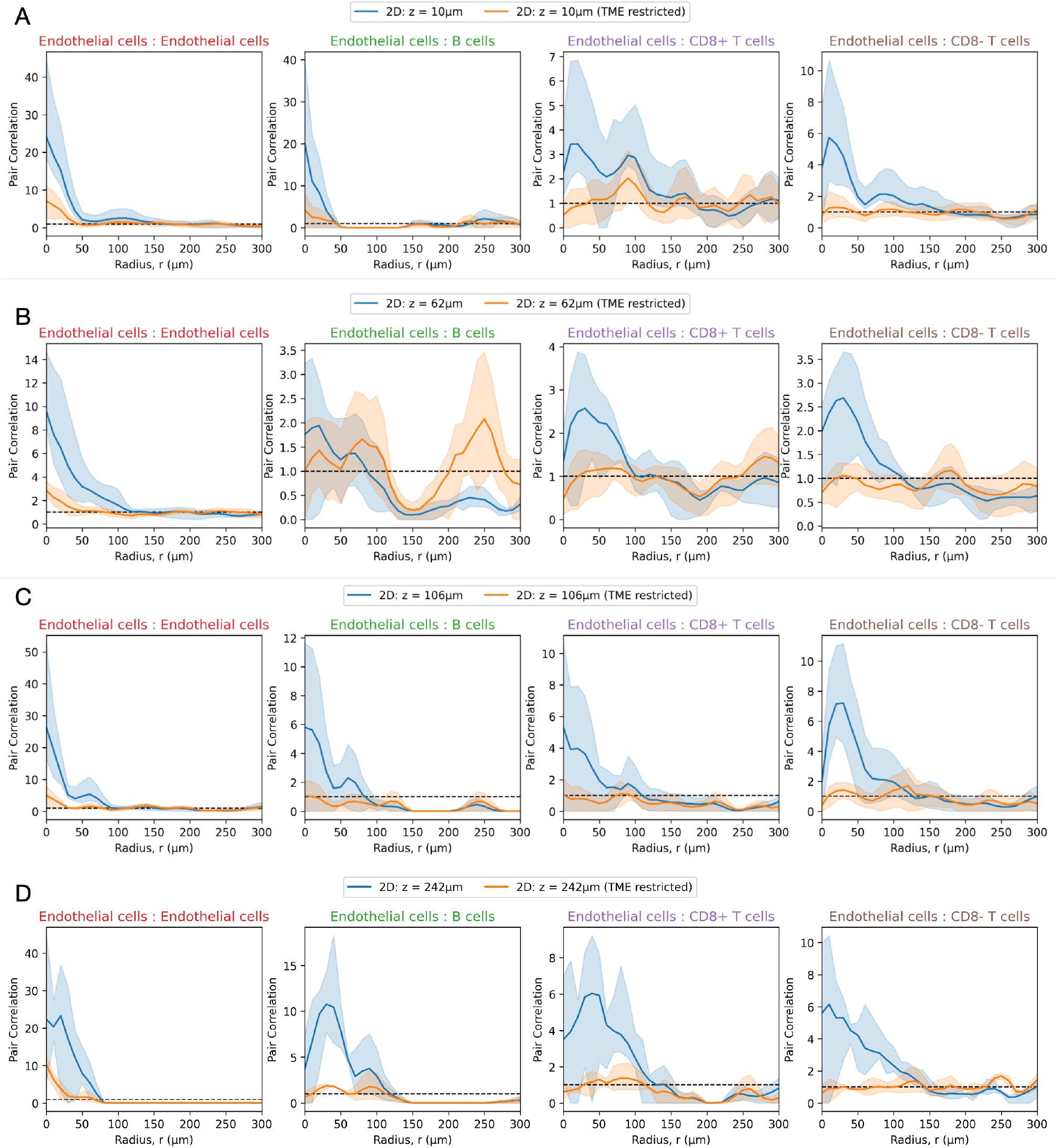
Endothelial–immune spatial associations across additional 2D serial sections of the breast carcinoma IMC sample. Cross-netPCF estimates between endothelial and immune cell populations computed on individual 2D serial sections of the breast carcinoma IMC dataset. Curves are shown for both the full 2D tissue domain and the tumour microenvironment (TME)-restricted regions. (A) Section at *z* = 10 µm. (B) Section at *z* = 62 µm. (C) Section at *z* = 106 µm. (D) Section at *z* = 242 µm. Shaded regions denote 95% confidence intervals.

### 1.10 Extended information mammalian embryo analysis

#### Surface network construction

To construct spatial domains for netPCF analysis, we used the surface masks of the processed Optical Projection Tomography (OPT) images at E9.5 and E11.5 [42]. For each developmental stage, we generated a spatial network over the embryonic surface using an upper edge-length filtration threshold of *ϵ* = 0.15 µm. Each node in the resulting network was assigned continuous Wnt gene expression marks by summing the expression signal within a 0.2 µm neighbourhood in 3D space. This produced node-level continuous marks for each Wnt gene across the reconstructed embryonic surface. To enable comparison with categorical marked point pattern analyses, continuous expression values were also converted into binary positive or negative expression labels using Otsu’s thresholding method [49]. Thus, for each embryo, we obtained a spatial network representation of the surface equipped with both continuous and categorical Wnt expression marks (Fig. 9A).

#### netPCF parameters

For all netPCF cross-netPCF and weighted-netPCF estimations, we used a maximum interaction radius of 3 mm, chosen to provide coverage across the majority of the reconstructed embryo surface. Kernel parameters were selected using the lower-bound bandwidth estimates derived in Supplementary Information 1.6. Specifically, we employed a kernel shape parameter of *n* = 2, corresponding to an Epanechnikov kernel, with bandwidth Δ*r* = 0.2 µm for all estimations. In weighted-netPCF estimates, the feature kernel exponent was set as *n* = 2, with a bandwidth of Δ*θ* = 0.1. These parameters provided sufficient kernel support over the reconstructed spatial network while maintaining stable correlation estimates.

As a validation of the underlying spatial network representation, the unmarked netPCF was computed over all network nodes. This yielded *g*^net^(*r*) ≈ 1 for all *r <* 3 mm, indicating approximate spatial homogeneity of the reconstructed point cloud under the network metric (Fig. 22). Consequently, the marked cross-netPCF signals observed in the biological analyses are attributable to the spatial organisation of cellular populations rather than to large-scale inhomogeneity in the underlying spatial point density.

**Fig 22.**
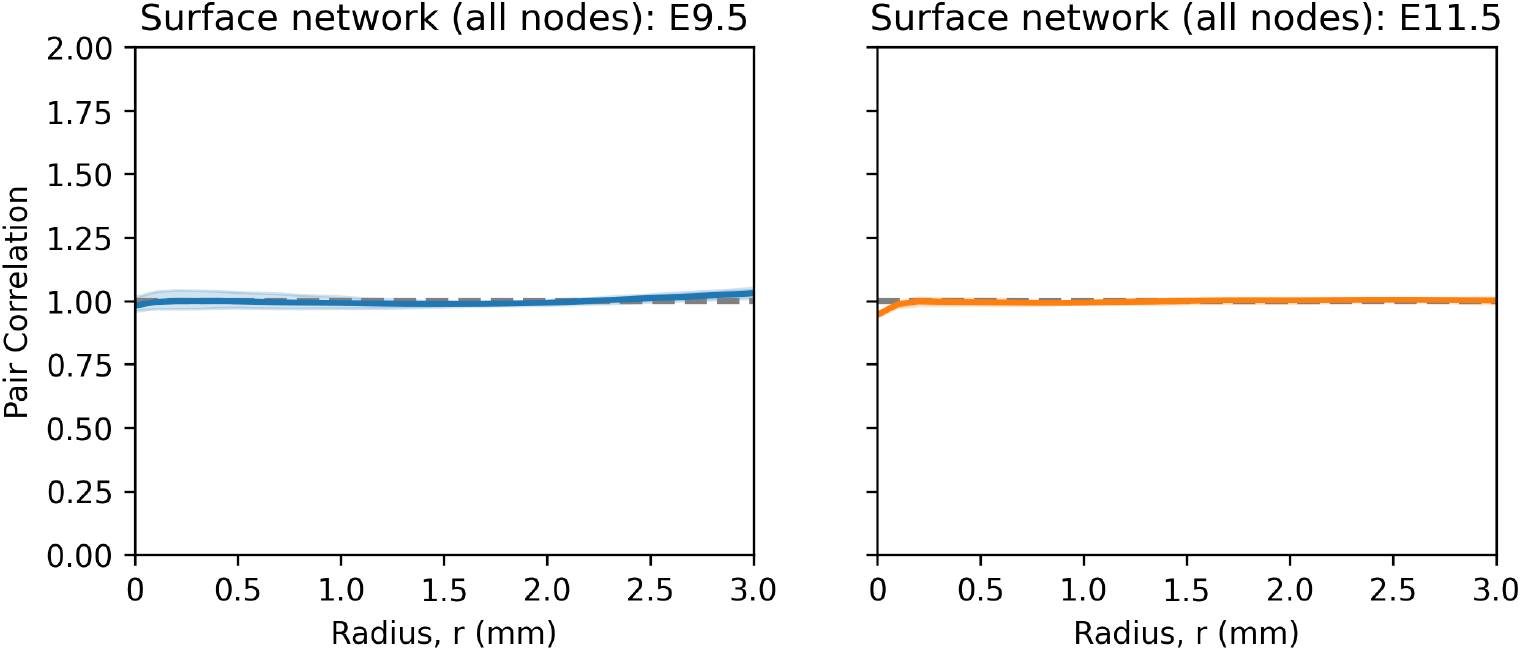
Pair correlation estimations of domain geometry: Mammalian embryos. netPCF estimates between all nodes of the murine embryo surface networks at E9.5 and E11.5. Shaded regions denote 95% confidence intervals.

